# Reduced flexibility in predictive tuning and contextual adaptation in autism: an EEG and behavioral study

**DOI:** 10.64898/2026.04.14.718519

**Authors:** Theo Vanneau, Seydanur Reisli, Chloe Brittenham, Michael J. Crosse, Sophie Molholm

**Author notes:** These authors contributed equally (Theo Vanneau, Seydanur Reisli).

## Abstract

The brain generates predictions to prepare for upcoming events. Because the environment is not perfectly predictable, the brain also estimates the certainty of these predictions and adjusts preparatory processes accordingly. Given that autistic individuals often resist even small changes to everyday routines, we hypothesized altered tuning of prediction certainty in autism. To test this, EEG was recorded from adolescents and young autistic adults (n = 20) and from age- and IQ-matched non-autistic adults (n = 19) during a probabilistic cued target identification task during which cue validity was systematically varied across four levels: 100%, 84%, 67%, and 33%. Participants were not informed of the cue-target validity nor when it changed. We focused on two neural signatures of anticipatory readiness, contingent negative variation (CNV) and alpha-band event-related desynchronization (α-ERD), and one of cognitive updating: the P3 to targets and to invalid (e.g., a non-target in place of the target) stimuli. Across groups, preparatory activity increased as contextual certainty decreased, with larger CNV amplitudes and stronger α-ERD preceding targets in lower-probability contexts, suggesting enhanced preparatory engagement under greater uncertainty. Furthermore, larger CNV amplitudes predicted faster reaction times, indicating functionally significant anticipatory dynamics. However, modulation of both neural preparation and response times as a function of cue-target probability was significantly reduced in the autistic group. In addition, autistic participants showed diminished probability-dependent modulation of the P3b to both targets and invalid stimuli, and coupling between anticipatory activity (CNV) and subsequent updating (P3b) was observed in non-autistic participants whereas it was absent in autism. Together, these findings suggest that while predictive mechanisms are present in autism, anticipatory processes are less flexibly tuned to contextual uncertainty and less effectively linked to subsequent cognitive updating. This reduced adaptability may reflect difficulty adjusting internal predictive models to changing environmental contingencies, potentially contributing to core features of autism such as resistance to change and insistence on sameness.

**Highlights:**

- Anticipatory brain mechanisms (CNV and alpha desynchronization) are present in autism and are behaviorally relevant, predicting faster responses.
- Autistic individuals exhibit reduced modulation of anticipatory CNV and alpha activity as a function of cue-target validity.
- P3b responses to both targets and invalid stimuli show diminished sensitivity to contextual probability in autism, consistent with altered prior updating.
- The link between anticipatory activity and cognitive updating (i.e., CNV to P3b) is disrupted in autism.
- P3a amplitude to invalid stimuli is reduced in autism, suggesting diminished engagement of violation-sensitive processes.
- Together, findings point to less flexible tuning of predictive mechanisms and reduced adaptation to contextual uncertainty in autism.

## Introduction

Far from being a passive mirror of reality that merely reacts to sensory input, the brain is now recognized as a predictive organ that continuously generates expectations about upcoming events. These predictions, shaped by prior experience and updated with incoming sensory evidence, guide perception and allow faster, more efficient processing by focusing resources on what is most likely to occur, while also reducing surprise and limiting the processing demands associated with unexpected events (Bar, 2007; Gregory, 1980; Hohwy, 2017). Contemporary theories propose that the brain maintains an internal model of the world that generates top–down predictions about forthcoming sensory input. Incoming signals are compared with these predictions, and discrepancies produce prediction errors that update the model (Bar et al., 2006; Nobre et al., 2007; Rao & Ballard, 1999). Crucially, both predictions and prediction errors are weighted by their estimated precision, reflecting the reliability of contextual information. Adaptive behavior therefore requires the flexible adjustment of predictions and their precision as environmental uncertainty changes (Friston & Kiebel, 2009).

Autism is characterized by impoverished social communication and restricted and repetitive behaviors (American Psychiatric, 2013) and is diagnosed in 1 out of 32 individuals in the United States and in 1 in 100 worldwide (Maenner et al., 2023; Zeidan et al., 2022). Despite extensive research, the neurobiological mechanisms underlying autism remain poorly understood. In recent years, predictive processing frameworks have gained traction in autism research (see Cannon et al., 2021 for a review), as they are considered to offer a unifying theoretical account capable of generating testable hypotheses and potentially explaining a wide range of sensory, cognitive, and motor characteristics associated with the condition (Gomot & Wicker, 2012; Van de Cruys et al., 2014). Within this framework, core features of autism have been interpreted as reflecting alterations in predictive inference. For example, social communication difficulties may arise from challenges in constructing generative models that allow reliable anticipation and interpretation of complex social cues (Chambon et al., 2017; Palmer et al., 2015). Likewise, repetitive behaviors and resistance to change may partly reflect attempts to restore predictability and thereby maintain a sense of stability or comfort in environments where violations of expectation are experienced as especially salient or disruptive (Gomot & Wicker, 2012). Although these accounts have generated considerable interest, empirical findings are equivocal (Cannon et al., 2021; Pesthy et al., 2023; Sapey-Triomphe et al., 2025; Shi et al., 2025), suggesting that predictive mechanisms may be selectively altered depending on the context (Sapey-Triomphe et al., 2021), the level of processing involved, or the specific computational stage under investigation.

Neurophysiological studies have examined predictive processing in autism primarily by assessing sensory responses to expected versus unexpected events, most commonly using oddball paradigms and mismatch negativity (MMN) measures (Sapey-Triomphe et al., 2025). Within predictive coding frameworks, discrepancies between incoming sensory input and prior expectations generate prediction errors. The existing literature suggests that autistic individuals may process these prediction errors differently, often showing reduced MMN amplitudes relative to non-autistic participants (Chen et al., 2020; Sapey-Triomphe et al., 2025; Schwartz et al., 2018). Several predictive coding accounts have been proposed to explain these altered neural responses to unexpected stimuli in autism. The hypo-prior hypothesis suggests that autistic individuals rely less strongly on prior expectations (Pellicano & Burr, 2012), which may result in weaker sensory predictions and, in turn, smaller MMN responses. By contrast, the HIPPEA account proposes that prediction errors are assigned excessively high and inflexible precision in autism (Van de Cruys et al., 2014). Although this framework could, in principle, predict enhanced responses to unexpected events, it also implies less stable learning of environmental regularities, which could ultimately diminish the distinction between standard and deviant stimuli. As a result, predictive coding theories do not converge on a single straightforward prediction regarding neural responses to unexpected stimulation but rather suggest effects that may vary depending on context. Furthermore, as highlighted by a recent meta-analysis (Sapey-Triomphe et al., 2025), MMN amplitude and latency alone often do not reliably distinguish autistic from non-autistic individuals, underscoring the need for more sensitive and nuanced paradigms to identify robust group differences.

Predictive processing involves not only the detection of mismatches once a stimulus has appeared, but also preparatory mechanisms that shape neural activity in advance of expected events. However, these anticipatory processes have received comparatively less attention. Predictions about task-relevant stimuli can modulate neural activity prior to stimulus onset, reflecting anticipatory allocation of attentional resources, changes in cortical excitability, and preparation for action (Bidet-Caulet et al., 2012; Thillay et al., 2016). From this perspective, predictive processing can be understood as a sequence of temporally organized operations that includes anticipatory preparation, sensory comparison between prediction and input, behavioral implementation, and subsequent model updating (Bidet-Caulet et al., 2012; Garrido et al., 2009; Rao & Ballard, 1999).

Among the limited studies examining neural mechanisms of anticipation in predictive contexts in autism, Thillay and colleagues (2016) found that anticipatory activity in autistic adults was less modulated by changes in contextual predictability (Thillay et al., 2016). Using a cued target paradigm and EEG recordings, they found that autistic participants exhibited neural signatures of anticipation both for predictable targets, presented after a cue sequence, and for unexpected targets presented without a preceding cue. Similarly, in a cued-target detection EEG study in autistic children, neural signatures of anticipation did not modulate by context (Beker et al., 2021). This pattern suggests a reduced ability to flexibly adjust anticipatory cortical activity according to the predictive value of contextual information, as if anticipatory resources were deployed more broadly, rather than selectively deployed toward meaningful events.

Importantly, predictive coding frameworks emphasize that the brain must continuously estimate the reliability or certainty of predictions, adjusting the gain applied to predictions and prediction errors and model updating accordingly (Keller & Mrsic-Flogel, 2018). In dynamic environments, this requires flexible modulation of anticipatory preparation and model updating as the reliability (and meaning) of contextual cues changes. However, relatively few studies have directly manipulated prediction certainty to examine how these neural mechanisms adapt across contexts in autism.

To address this question, we developed a probabilistic cued target detection task in which cue validity was systematically manipulated (100%, 84%, 67%, and 33%), leading to different contextual certainty without modification of the perceptual information. Participants were first exposed to a fully stable context (100% validity) before encountering blocks with different levels of contextual certainty (as manipulated by cue-validity). Using EEG, we examined how autistic individuals and matched non-autistic controls process these varying levels of contextual certainty. We focused on three neural markers potentially involved in predictive processing and model updating that are sensitive to target probability: the Contingent Negative Variation (CNV), a slow fronto-central potential reflecting anticipatory preparation (Kononowicz & Penney, 2016; Tecce, 1972; Walter et al., 1964); posterior alpha-band event-related desynchronization (α-ERD), indexing cortical excitability and attentional allocation to predicted targets (Boettcher et al., 2020; Hanslmayr et al., 2011; Klimesch et al., 2007; Pfurtscheller & Klimesch, 1992; Rohenkohl & Nobre, 2011), and the P3b, a centro-parietally focused positive going response, associated with internal model updating (Bidet-Caulet et al., 2012; Polich, 2007; Polich, 2011).

Our goal was to determine whether predictive processing alterations in autism reflect globally reduced anticipatory and updating mechanisms, or rather a reduced capacity to flexibly modulate these mechanisms as contextual certainty changes. We first examined whether canonical neural signatures of anticipatory preparation (alpha ERD and CNV) and model updating (P3b responses) were present and behaviorally relevant in autistic participants. We then tested whether these neural responses scale with target probability, as typically observed in non-autistic individuals. Finally, we investigated how anticipatory processes, model updating, and behavioral responses are coordinated by examining the relationships between alpha-band desynchronization, CNV amplitude, P3b modulation, and reaction times. We expected non-autistic participants to show graded modulation of these neural markers as cue validity increased, reflecting flexible tuning of prediction certainty and response readiness. In contrast, we hypothesized that autistic participants would show reduced modulation of these neural signatures across probability levels and altered coordination between neural dynamics and behavior.

## Methods

### Participants

The initial sample comprised 23 non-autistic adults and 23 autistic adults. Following exclusions due to EEG data quality and behavioral response criteria (see below), the final analyzed sample included 19 non-autistic (17-28 years old; 20.8 ± 2.6; mean ± standard deviation) and 20 autistic (17-28 years old; 19.7 ± 2.49) individuals. Autism diagnoses were confirmed using the Autism Diagnostic Observation Schedule, Second Edition (ADOS-2; Lord et al., 1994), and expert clinical judgment by a licensed psychologist at the Human Clinical Phenotyping Core of the Rose F. Kennedy Intellectual and Developmental Disability Research Center at the Albert Einstein College of Medicine. Participants were recruited without regard to sex, race, or ethnicity. Exclusionary criteria for both groups included a performance IQ below 80, a history of head trauma, premature birth, a current psychiatric diagnosis, or a known genetic syndrome associated with a neurodevelopmental or neuropsychiatric condition. Attention deficit/hyperactivity disorder (ADD/ADHD) was not used as an exclusion criterion for the autism group, given its high comorbidity with autism (Rong et al., 2021). Exclusion criteria for the non-autistic group additionally included a history of developmental, psychiatric, or learning difficulties, and having a first-degree relative with an autism diagnosis. Participants who were on stimulant medications were asked to not take them at least 24 hours prior to the experiment. All procedures were approved by the Institutional Review Board of the Albert Einstein College of Medicine and adhered to the ethical standards outlined in the Declaration of Helsinki. All participants either consented or assented to the procedures. For participants under the age of 18, a parent or guardian signed an informed consent. The protocol was approved by Institutional Review Board of the Albert Einstein College of Medicine. Participants received nominal recompense for their participation (at $15 per hour). IQ was measured via the Wechsler Abbreviated Scale of Intelligence (Simard et al., 2015).

### Experimental Procedure

We designed a task to probe the ability to adjust predictions based on changing probabilities (certainty) in the environment.

#### Stimuli

Visual stimuli were presented to the participant, one at a time, on a computer screen (Dell UltraSharp 1704FPT) at a viewing distance of 65 cm in an electrically shielded room (International Acoustics Company, Bronx, New York). Stimuli consisted of basic shapes presented in gray on a black background for 100ms, with an 850ms inter-stimulus interval (ISI). Participants performed a target detection task in which they responded as quickly as possible to the final item of a *target-sequence*. A target-sequence was either three arrows, the first upward-facing, the second rightward-facing, and the final downward-facing, or three parallelograms, the first left-tilted, the second straight, and the final right-tilted. The stimuli in these sequences are referred to as *Cue1*, *Cue2*, and *Target* (Fig. 1A). When sequences were not completed with a target, a circle, diamond, or triangle shape was presented instead, which we refer to as an *invalid item*. These shapes were also used as *fillers*, presented once or twice after invalid items or targets. To ensure that participants were responding to the shape sequence and not just the final shape in the sequence, *catch* trials in which the final shape was presented after filler shapes were also included.

**Figure 1.**
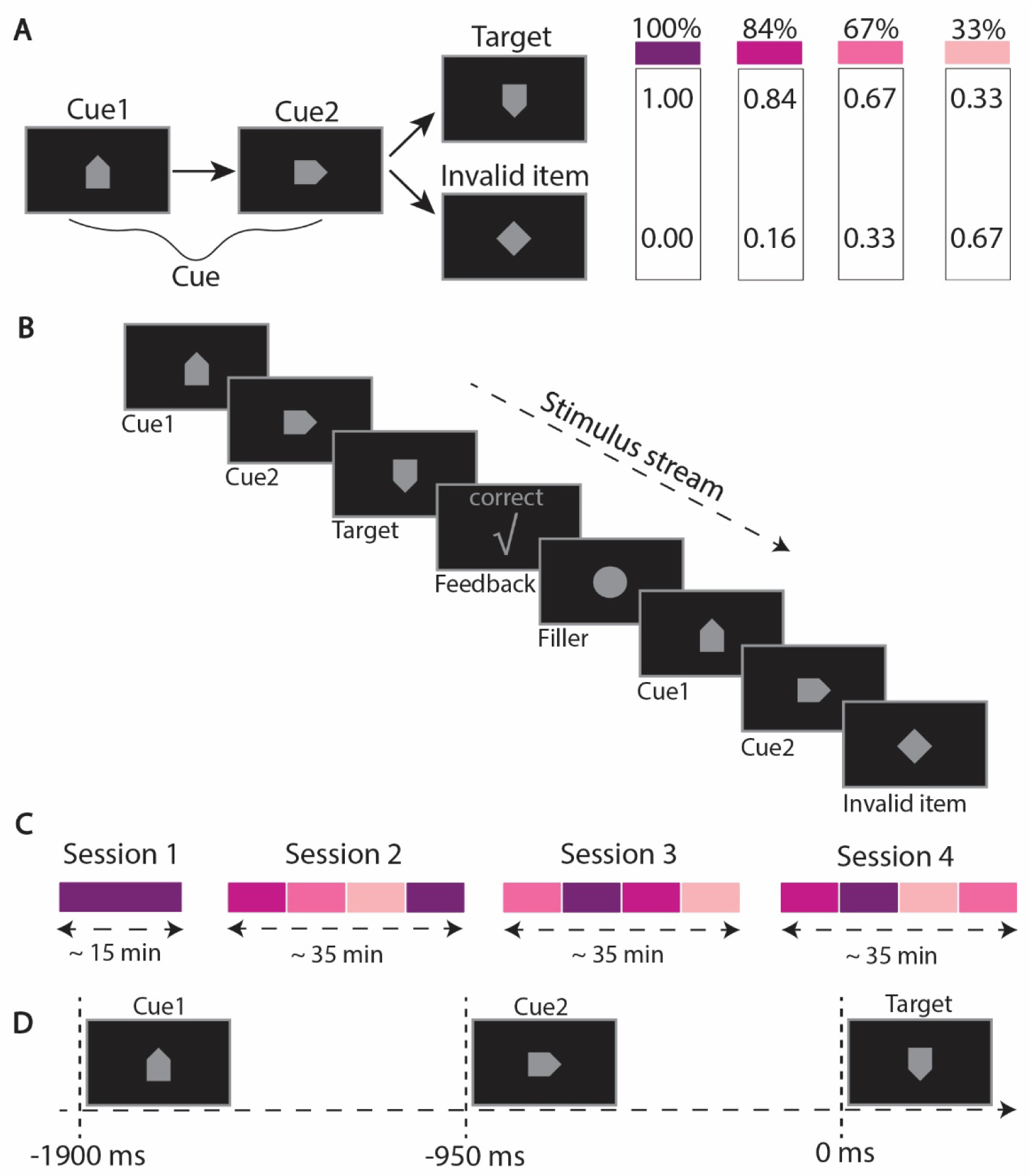
The Sequential Probabilistic Task. **(A)** Participants responded to target sequences of stimuli while the probability of sequence completion was manipulated at four levels. Stimuli consisted of basic shapes presented sequentially to the participant. There were two possible target sequences (only one is represented here): A sequence of 3 different arrows or 3 different parallelogram shapes were presented in specific orders. The participant’s task was to respond after sequence completion with a mouse click while withholding the response when the sequence was completed with an invalid item. There were 4 different probabilities of having a target completing the pattern initiation. **(B)** A sample stimulus stream. The subject responds with a mouse click after completion of a three-item target pattern, followed by a feedback message appearing on the screen. **(C)** The order of cue validity conditions throughout the experiment is shown for a sample participant (See methods for blocks details). **(D)** Illustration of how a target sequence unfolds over time relative to EEG epochs, with time 0 placed at target onset.

#### Cue Validity Conditions

Throughout the experiment, the probability that a target-sequence was completed varied across four levels, in ∼10-minute blocks (Fig. 1C). Pattern initiations, always represented by Cue1 of the pattern followed by Cue2, were completed with the target stimulus 100%, 84%, 67% or 33% of the time, comprising four cue validity conditions (Fig. 1A). Participants were not informed of the cue validity condition they were in or when it changed. Two target-sequences (arrows or parallelograms) were used with the goal of making the task more engaging. They were presented with equal probability within a given cue validity condition.

#### Blocks

Stimuli were presented in mini blocks of ∼1.5 minutes, separated by pauses during which time participants could rest. Each mini block was composed of 24 pattern initiations (Cue1 followed by Cue2). Pattern initiations were completed with the target 24 (100%), 20 (83%), 16 (67%), or 8 (33%) times depending on the cue validity condition. Participants pressed the mouse key to initiate the next mini block. Blocks of a given cue validity condition were composed of between 4 and 6 mini-blocks.

#### Remote Familiarization

To briefly familiarize participants with the stimuli and task prior to the experiment, they performed the task remotely (100% cue validity condition only) for six minutes using the Neurobehavioral Systems mobile app on their smart phone or tablet, one day before the experiment. Completion of the training was verified before participants attended the laboratory session.

**Instructions Part 1**. The following instructions were printed on the screen in four parts, both for remote familiarization and the first experimental session:

*“You will see a shape in the middle of the screen. The shape will change about every second. Sometimes 3 consecutive shapes appear in the orders below, which we call a pattern. There are two target patterns: (pattern shapes were shown to the participant below this sentence). Your job is to touch the screen (or press the mouse button) after Pattern 1 or Pattern 2 is completed. Try to be both quick and accurate. Remember, you should respond after the pattern is completed. You can ignore any other shape. Let’s practice!”*

### Experiment Sessions

The experiment was composed of four sessions performed on a single day, separated by 15 to 30-minute breaks (Fig. 1C). In Sessions 1 and 2, the cue validity conditions were presented in the same order to all participants, whereas in Sessions 3 and 4, cue validity condition order was pseudo-randomized. Session 1 consisted of 7 mini-blocks of 100% cue validity condition. In Session 2, conditions were presented in the order of 84%, 67%, 33%, and 100%. Participants usually took a lunch break after Session 2, and a ∼15-minute break between Session 3 and Session 4. In Sessions 3 and 4, cue validity conditions were presented in a pseudo-randomized order (sample order is shown in Fig. 1B).

**Instructions Part 2**. At the end of the first session, participants were informed that going forward, the cues would not always be followed by the target, and that in these cases they should withhold their response.

**Feedback**. For both remote and in-lab tasks. To keep the participant on-task, visual feedback was provided: “correct” for responses to targets that fell within the response window of 100 to 950 ms; “miss” if they did not respond within 950 ms of the target; “too early” for responses occurring within 100 ms of target presentation (assumed to be anticipatory); and “wrong” for responses to a non-target. Feedback text was accompanied by an icon (a “✓” for correct, “x” for wrong, “!” for miss or too early). The feedback stimulus was presented for 200 ms.

### EEG Recordings & Preprocessing

EEG data were recorded at a sampling rate of 512Hz using 160 channels BioSemi Active II system (using the CMS/DRL referencing system) with an anti-aliasing filter (−3 dB at 3.6 kHz). Analyses were conducted in Python (version 3.11) using MNE (Gramfort et al., 2013) and custom scripts are available at https://github.com/tvanneau/SPLT. Bad channel detection was performed using the function NoisyChannels (with RANSAC) from the pyprep toolbox (Bigdely-Shamlo et al., 2015). If more than 15% of the channels were detected as bad, the participant was rejected (4 non-autistic and 2 autistic). Bad channels were interpolated using spline interpolation (Perrin et al., 1989). EEG was filtered using a FIR band-pass filter (0.01-40Hz), and Independent Component Analysis (ICA) on 1Hz high-pass EEG was used to identify and manually reject eye-related components (blinks/saccades). Epochs were created from −2200 to +800ms around each imperative stimulus (i.e., valid and invalid targets). Baseline correction was applied using the −2000 to −1900 ms interval, corresponding to the 100 ms preceding *Cue1*, in order to ensure a stimulus-free baseline. For all subsequent analyses, EEG data were re-referenced to the average of all electrodes.

### Data Analysis

For analyses of the behavioral and EEG data, we excluded the first session, which consisted of a 15-minute block with 100% target probability. This initial session served to familiarize participants with the task with the additional goal that they would build a high certainty predictive model for the starting point of the experiment. Therefore, only data from sessions 2, 3, and 4 were included in the analysis.

### Behavioral data analysis

For each trial, we analyzed participants’ responses by measuring reaction time (RT) when a button press occurred, either in response to a target (classified as a hit) or to an invalid item (classified as false alarm). A miss was recorded when no response followed a target. For each condition and participant, we calculated: Mean reaction time for targets, mean reaction time for invalid items, hit rate (proportion of targets correctly detected), false alarm rate (proportion of responses to invalid items), and number of misses (no response to targets). If the hit rate of a participant was below 50% in the 100% target probability condition, the participant was excluded from further analysis (1 autistic participant).

### ERP data analysis

ERP analyses combined theory-driven region-of-interest (ROI) approaches with complementary cluster-based permutation tests. For components expected to scale parametrically with cue–target probability (four levels: 100%, 84%, 67%, 33%), modulation was quantified using within-subject linear slopes. For responses to invalid stimuli (three probability levels), linear mixed-Xeffects models (LMMs) were used instead of slope estimation.

### Anticipatory Activity: CNV

To examine anticipatory brain activity, we focused on the contingent negative variation (CNV), a slow negative ERP component developing in anticipation of an expected event. Based on prior literature (Kononowicz & Penney, 2016; Tecce, 1972; Walter et al., 1964) and topographical maps (see Figure 2C), we selected a region of interest (ROI) comprising 18 central electrodes around Cz (Figure 2A), consistent with the canonical central CNV distribution reported previously (Galvao-Carmona et al., 2014; Michelini et al., 2018; Wang et al., 2020). CNV amplitude was quantified as the mean voltage during the 50ms immediately preceding target onset (−50 to 0ms). This late window captures the final build-up of preparatory activity and is particularly sensitive to temporal expectancy and event probability (Bauer et al., 1992; Korunka et al., 1993). Given the graded probability manipulation, focusing on the terminal preparatory phase allowed us to assess the anticipatory state immediately before stimulus presentation. To confirm the presence of a CNV, mean amplitudes in the Pre-Target window (−50 to 0ms) were compared to a control window preceding Cue 2 (Pre-Cue, 50ms) using an LMM with Time-Window and Group as fixed effects and Subject as a random intercept, as described in the data analysis section. All probability conditions were pooled for this analysis. To quantify contextual modulation, CNV slopes were computed for each participant by regressing normalized CNV amplitudes onto target probability (coded 1.00, 0.84, 0.67, 0.33). Prior to slope estimation, amplitudes were mean-centered within participant to isolate relative probability-driven modulation from overall amplitude differences. Described in more detail in the data analysis section, complementary time-cluster permutation tests were performed across the full pre-target interval to capture effects outside the predefined window.

**Figure 2.**
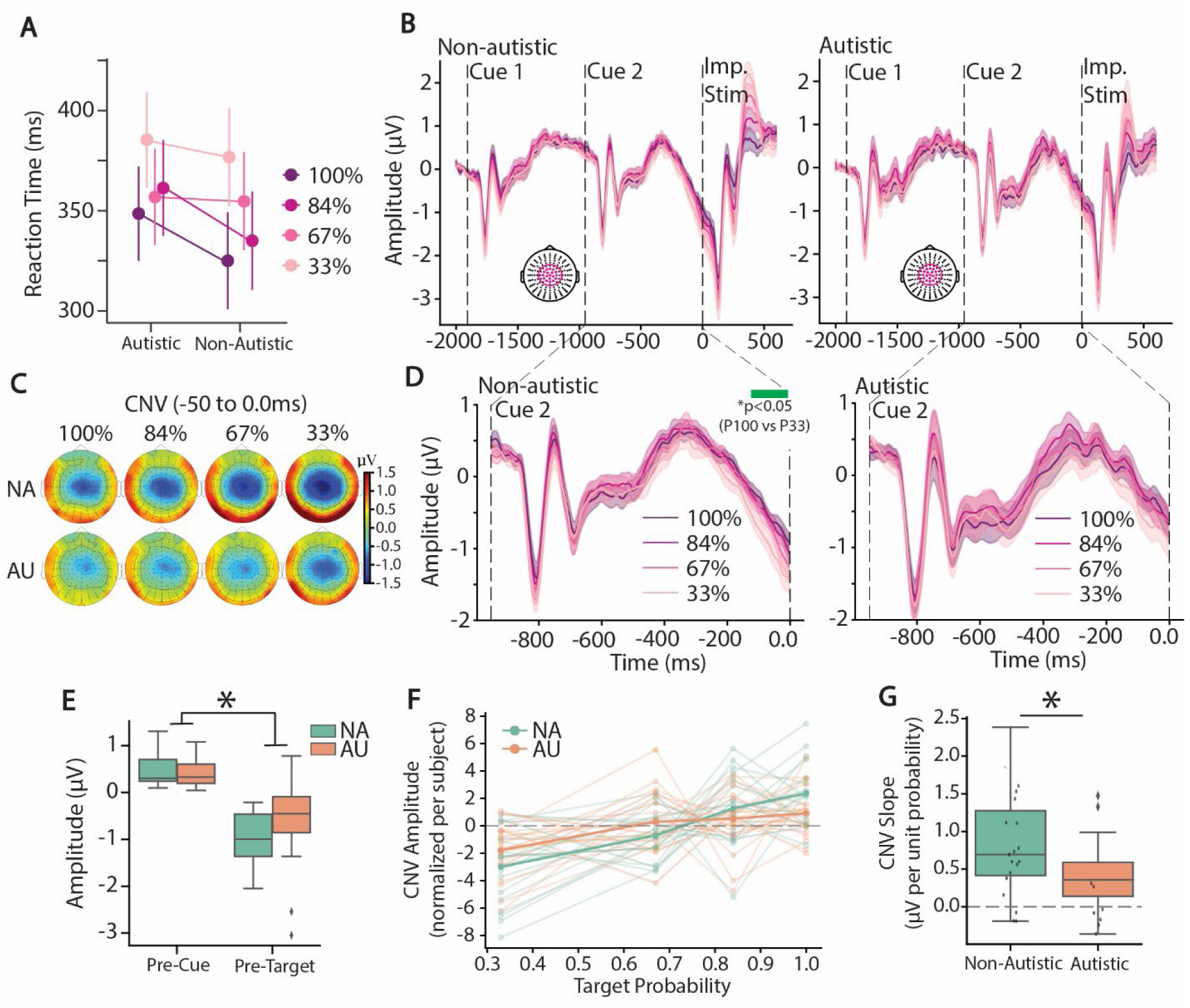
Reduced modulation of the CNV by target probability in autistic group. **(A)** Reaction times for 100% (violet line), 84% (light violet), 67% (pink) and 33% target probability block (light pink), for non-autistic (NA) and autistic (AU) groups. **(B)** Event-related potentials (ERPs) averaged over a cluster of central channels over the full sequence with Cue 1 (first dashed line, −1900ms), Cue 2 (second dashed line, −950ms) and both imperative stimuli (target and invalid target) at 0ms, for 100% (violet line), 84% (light violet), 67% (pink) and 33% target probability block (light pink), for non-autistic on the left and autistic on the right. **(C)** Topographical representation of the 50ms before imperative stimulus onset for each target probability from 100% on the left to 33% on the right, for non-autistic (top row) and autistic (bottom row). **(D)** Focus on the CNV timing (between Cue 2 and imperative stimulus onset). Blue rectangle illustrates significant statistical differences assessed with cluster-based permutations (α = 0.05). (**E**) Average amplitude for the 50ms before Cue 2 (Pre-Cue) and for the 50ms before imperative stimulus onset (Pre-Target) over the cluster of central channels selected for the ERPs for non-autistic (green) and autistic (orange) averaged between the 4 targets probability conditions. * Indicates significant differences assessed by LMMs. **(F)** Linear regression of the CNV slope by target probability with group-mean curve for non-autistic (blue) and autistic (orange) as individual regression curve in corresponding color. **(G)** Slope of the CNV regression curve for each group. * Indicates significant differences assessed by ANOVA.

### Anticipatory Activity: Alpha power

Time–frequency analyses were performed using complex Morlet wavelet convolution (MNE-Python tfr_morlet) from 4–20 Hz (100 frequency steps, FWHM = 2 Hz; 3 cycles at 4 Hz to 10 cycles at 20 Hz). This provided a temporal resolution of ∼281 ms at 4 Hz and ∼187 ms at 20 Hz (Cohen, 2014). To isolate induced activity, the ERP was subtracted from each trial prior to decomposition (Kalcher & Pfurtscheller, 1995). Power values were then baseline-normalized (percent change) relative to the 100ms window preceding Cue 1 (from −2000 to −1900ms relative to target onset). Analyses focused on alpha-band activity (7–13 Hz) averaged across 14 parieto-occipital electrodes (Figure 4A), consistent with prior anticipatory alpha studies (Boettcher et al., 2020; Gould et al., 2011; Li et al., 2023; Tarasi et al., 2022). Mean alpha power was extracted from −100 to 0ms before target onset, capturing late-stage preparatory modulation of cortical excitability (Gould et al., 2011; Rohenkohl & Nobre, 2011). Alpha slopes were computed using the same within-subject normalization and linear regression procedure applied for measurement of the CNV. Finally, we computed the alpha power difference between the highest (100%) and lowest (33%) probability conditions. Both the condition-wise alpha time course and the corresponding time–frequency representations are shown in Figure 6.

### Target-Evoked Updating: P3b

To assess cognitive updating, we analyzed the P3b component elicited by targets (Polich, 2007; Squires et al., 1975). Inspection of the grand-average topographical maps (Figure 6B) revealed a canonical centro-parietal distribution. Based on this, we defined a region of interest (ROI) comprising 12 centro-parietal electrodes (Figure 6A), consistent with the typical P3b topography reported in prior literature (He et al., 2001; Polich, 2007; Polich, 2011; Smith et al., 1990). Because epochs were baseline-corrected prior to Cue 1, the CNV was present at stimulus onset and could inflate the amplitude differences for the P3b. To avoid confounding baseline shifts, no additional pre-target baseline correction was applied. Instead, P3b amplitude was quantified using a trough-to-peak method: the most negative voltage within 130–200ms post-target onset was subtracted from the most positive voltage within 300–350ms. This approach minimizes contamination from slow baseline shifts and provides a robust amplitude estimate. As with the CNV, P3b slopes were computed by regressing within-subject normalized amplitudes onto target probability.

### Responses to Invalid Stimuli: N2, P3a, and P3b

To examine responses to expectancy violations, analyses focused on invalid trials in which participants correctly withheld responses. This approach parallels correct-rejection trials in go/no-go paradigms. The N2 was quantified over a fronto-central ROI based on observed topography (Figure 7A) and prior literature (Bokura et al., 2001; Folstein & Van Petten, 2008). Mean amplitude was extracted from 260–280ms, centered on the grand-average minimum. The P3a was measured over the same central ROI from 380–400ms, corresponding to the fronto-central positivity following the N2 (Bokura et al., 2001; Polich, 2011). Concerning the P3b, compared to the valid trials, a later window (460–480ms) was used. This shift was motivated by (i) prior evidence that expectancy violations produce delayed P3b peaks relative to valid targets (Lasaponara et al., 2018; Schuller et al., 2025), (ii) Examination of the waveforms (Fug X), and (iii) spatio-temporal cluster permutation results revealing group differences in this later range (Figure 7G). Because only three invalid-probability levels were available (16%, 33%, 67%), slope estimation was not performed. Instead, LMMs were fitted with Component Amplitude as the dependent variable and Group and Condition as fixed effects, with Subject as a random intercept.

### Statistical Analysis

Statistical analyses, described in detail below, were conducted using Jamovi (version 2.3.28) for Analysis of Variance (ANOVA) and linear mixed-effects models (LMMs) and in MNE-Python for permutation-based testing (Maris & Oostenveld, 2007).

#### Behavioral Data

Reaction times (RTs) were compared across groups and cue validity conditions using LMMs to account for within-subject variability. Age was included as a covariate, and all fixed effects and their interactions were tested. We report F-values from the omnibus test of fixed effects, along with partial eta squared (η²ₚ) as the effect size and associated p-values (α = 0.05). The Shapiro–Wilk test was used to confirmX the normality of residuals.

#### EEG Data

Temporal statistical comparisons of EEG measures averaged over predefined electrode clusters (e.g., ERP amplitudes or alpha power) were performed using temporal cluster-based permutation tests (Maris & Oostenveld, 2007), implemented via the permutation_cluster_test function in MNE (1,000 permutations, two-sided F-test, α = 0.05). This method compares observed cluster-level statistics (e.g., maximum t-values) against a null distribution generated by randomly permuting the data. Additionally, spatio-temporal cluster-based permutation tests were applied to the difference wave between the 100% and 33% conditions (Figure 3A for the pre-target window; Figure 7G for the brain response to invalid stimulus) using MNE’s spatio_temporal_cluster_1samp_test function (1,000 permutations, two-sided one-sample t-test, α = 0.05). Group differences in the slope of modulation of the CNV, P300, and alpha power across target probability levels were tested using one-way ANOVAs (α = 0.05). Concerning the average amplitude for the N2, P3a and P3b in response to the invalid stimulus, we performed linear mixed models with the average amplitude of the corresponding component as the dependent variable and Group (NA and AU) and Condition (16, 33, 67% of invalid probability) as fixed factor, with Subject as a random effect to account for repeated measures.

**Figure 3.**
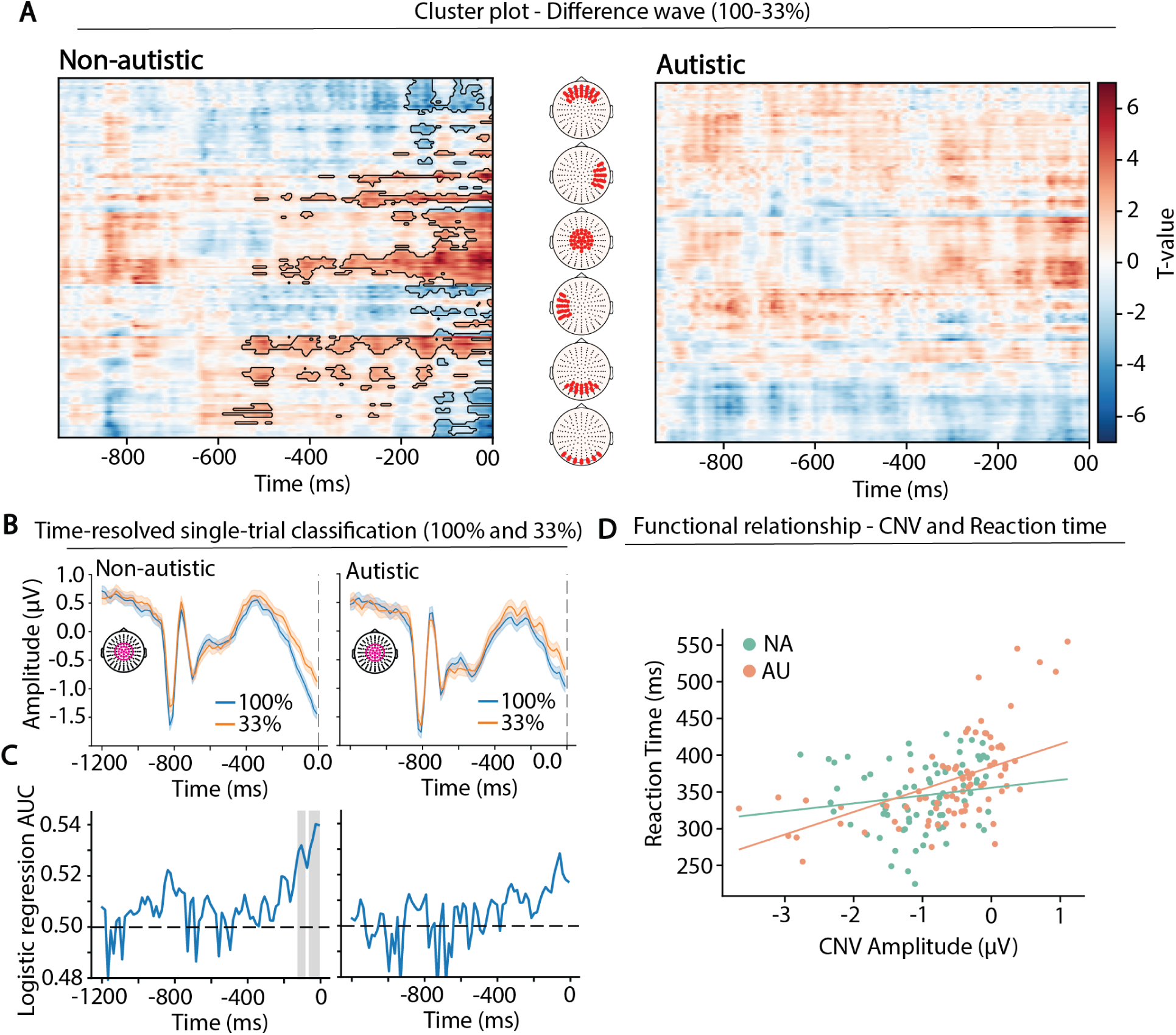
Reduced CNV modulation by target probability in Autistic group but maintained functional link to behavior. (**A**) Spatio-temporal cluster plots showing the t-values (one-sample t-test) of the 33–100% difference wave at each time point and electrode for non-autistic (left) and autistic (right) participants. Significant clusters are outlined in black (cluster-based permutation test). (**B**) Grand-averaged single-trial waveforms from a central electrode cluster for the 100% and 33% conditions in non-autistic (left) and autistic (right). Below: time-resolved classification accuracy (AUC of ROC curve) distinguishing 100% vs 33% trials using a leave-one-subject-out approach. Time points with above-chance classification are highlighted by gray rectangles (cluster-based permutation statistics). (**D**) Scatterplots showing the relationship between CNV amplitude and reaction time for non-autistic (blue) and autistic (red). Points represent condition-wise averages per participant. Solid lines depict linear mixed-effects model predictions (random intercept for Subject), controlling for target probability (Condition).

#### Linking anticipatory signals to behavior and model updating (condition-wise LMMs)

Analyses were conducted on condition-wise averages for each participant (four cue-validity levels per individual). Linear mixed-effects models (LMMs) were fitted with a random intercept for Subject to account for repeated measures across conditions. To test behavioral relevance, reaction time (RT) was modeled as a function of CNV amplitude (and, in a separate model, induced alpha power), including Condition as a fixed effect to account for systematic differences in cue–target probability across blocks. Group and its interaction with the neural predictor (e.g., CNV × Group) were included to test whether brain–behavior coupling differed between autistic and non-autistic participants. To assess coordination between anticipatory signals, CNV amplitude was modeled as a function of induced alpha power, again including Condition and testing for Group interactions. Although both measures were extracted from the late preparatory interval preceding target onset (CNV: −50 to 0ms; alpha: −100 to 0ms), they reflect partially distinct neural processes, as evidenced by their differing scalp topographies and spectral characteristics. Importantly, alpha estimates were derived from non-phase-locked (induced) activity following ERP subtraction, ensuring that the CNV-related evoked response did not contribute to the alpha measure. To examine whether anticipatory activity predicted subsequent cognitive updating, P3b amplitude was modeled as a function of CNV (and, separately, induced alpha power), controlling for Condition and testing Group × predictor interactions. Predictors were mean-centered to facilitate interpretability; fixed-effect slopes (β) quantify associations after adjusting for Condition and within-participant dependence.

#### Time-resolved Single-trial Classification

To assess whether CNV amplitude could discriminate target probability conditions at the single-trial level, we performed a time-resolved decoding analysis using a leave-one-subject-out (LOSO) cross-validation scheme. At each time-point, a logistic regression classifier (L2 regularization, lbfgs solver) was trained to distinguish between high-probability (target_P100) and low-probability (target_P33) trials based on the amplitude of the EEG averaged of the cluster of channels corresponding to the CNV. For each iteration, data from all participants except one were used for training, and the left-out participant served as the test set. Features were z-scored within the training set and applied to the test set. Classification performance was quantified using the area under the receiver operating characteristic (ROC) curve (area under the curve; AUC), averaged across folds. To assess significance while correcting for multiple comparisons across time, we applied a cluster-based permutation approach. At each time point, AUC values above a threshold of 0.525 (i.e., 0.025 above chance) were grouped into clusters. The cluster statistics were computed as the sum of AUC differences from chance. We then generated a null distribution of the maximum cluster statistic by repeating the permutation procedure (1000 iterations) with shuffled condition labels within participants (Groppe et al., 2011; Maris & Oostenveld, 2007). Observed clusters were considered significant if their cluster statistic exceeded the 95th percentile of the null distribution (p < 0.05).

### Results Reduced contextual modulation of RTs and CNV in autism

#### Reaction times

A linear mixed-effects model revealed a strong main effect of Condition, *F*(3,111)=31.39, η²ₚ=0.45, *p*<.001, no main effect of Group, *F*(1,36)=0.90, η²ₚ=0.02, *p*=.34, and a significant Group × Condition interaction, *F*(3,111)=3.00, η²ₚ=0.07, *p*=.03. Follow-up comparisons indicated that non-autistic participants showed graded modulation of reaction times across probability levels, with slower responses at 67% compared to 100% (*p*<.001; see Supplementary Table 1). In contrast, autistic participants did not differ in reaction times across the 100%, 84%, and 67% conditions (all *ps* = 1.0).

#### Contingent Negative Variation

Visual inspection of the full ERP sequence, baselined prior to Cue1, revealed similar waveform morphology across NA and AU groups. All trial types (including both targets and invalid items) were pooled across conditions, yielding 280 trials on average per condition per participant. Both groups showed clear P1/N1s in response to each visual stimulation (Figure 2A). During the anticipatory interval between Cue2 and target onset (−950 to 0ms), a sustained negative-going shift was evident in both groups, consistent with a Contingent Negative Variation (CNV; Figure 2B). To formally verify the presence of the CNV, mean amplitudes in the 50ms preceding target onset (Pre-Target window) were compared with those in the 50ms preceding Cue 2 (Pre-Cue window). Averaged across probability conditions, a linear mixed-effects model revealed a robust main effect of time window (F(1,74) = 88.95, η²ₚ = 0.54, p < 0.001), confirming the presence of anticipatory activity across participants (Figure 2D). There was no significant main effect of group (F(1,74) = 1.86, η²ₚ = 0.025, p = 0.17), or group × window interaction (F(1,74) = 2.99, η²ₚ = 0.04, p = 0.08), indicating that both groups exhibited comparable overall CNVs.

We next examined whether CNV amplitude scaled with target probability. Visual inspection suggested stronger probability-dependent modulation in the control group (Figure 2B). This was confirmed by a cluster-based permutation analysis, which revealed a significant difference between the 100% and 33% conditions in NA participants during the late pre-target interval (most prominent from −135 to 0ms; Figure 2B, green rectangle), but not in the AU group. Topographical maps further showed that NA participants exhibited a progressively larger central negativity under lower target probability, consistent with increased preparatory engagement. In contrast, AU participants showed relatively little variation across conditions (Figure 2C). Group comparison revealed significantly reduced slope values in AU relative to NA participants (F(1,31) = 4.48, p = 0.042; Figure 2F). Importantly, slopes were significantly different from zero in both groups (NA: t = 5.04, p < 0.001; AU: t = 3.71, p = 0.001), indicating preserved anticipatory processing in autism but attenuated sensitivity to contextual probability.

### Reduced Sensitivity of the CNV to Probability at Spatiotemporal and Single-Trial Levels in Autism

We next tested whether CNV modulation differed between the two extreme cue-validity conditions (100% vs. 33% target probability). To avoid bias toward predefined electrodes or time windows, we conducted a spatio-temporal cluster-based permutation analysis on the 33–100% difference wave separately for each group (Figure 3A). In the non-autistic (NA) group, significant clusters emerged over central electrodes from approximately −200 to 0 ms before target onset, confirming robust modulation of anticipatory activity as a function of target probability. In contrast, no significant clusters were observed in the autistic (AU) group, indicating a lack of reliable context-driven modulation at the distributed spatiotemporal level.

To determine whether this reduced sensitivity was also evident at the single-trial level, we performed a time-resolved decoding analysis using leave-one-subject-out logistic regression (see Methods). In the NA group, classification accuracy exceeded chance beginning approximately 130 ms before target onset, indicating that trial-by-trial CNV amplitude reliably encoded target probability (100 vs 33%). In contrast, decoding performance in the AU group remained at chance throughout the anticipatory interval (Figure 3B). Thus, although a CNV was present in both groups at the evoked level, its capacity to discriminate contextual probability (both in distributed spatiotemporal patterns and at the single-trial level) was markedly reduced in autism.

Finally, we examined the behavioral relevance of the CNV by modeling reaction time as a function of CNV amplitude, including target probability as a covariate and testing for group differences. The linear mixed-effects model revealed a significant main effect of CNV amplitude (β = 30.7 ms per µV, SE = 7.9, p < .001), such that more negative CNV amplitudes predicted faster responses. The CNV × Group interaction approached significance (β = −20.1 ms, SE = 10.7, p = .06), reflecting a numerically stronger CNV–reaction time coupling in the autistic group (≈31 ms per µV) compared to the non-autistic group (≈11 ms per µV). These results demonstrate that anticipatory neural activity remains functionally linked to behavior in both groups, despite reduced contextual modulation in autism.

### Modulation of Induced Alpha-Band Activity by Target Probability

We next examined whether induced alpha-band activity indexed anticipatory preparation and whether it was modulated by target probability. Alpha power is widely interpreted as an inverse marker of cortical excitability, with alpha desynchronization reflecting increased neural engagement (Hanslmayr et al., 2011; Klimesch et al., 2007; Pfurtscheller & Klimesch, 1992). Prior studies have shown that alpha power typically decreases in anticipation of expected stimuli, consistent with preparatory sensory facilitation (Boettcher et al., 2020; Rohenkohl & Nobre, 2011; Thillay et al., 2016).

To first establish the presence of anticipatory alpha suppression, we compared occipital alpha power during the 100ms preceding Cue 2 with the 100ms preceding target onset. A linear mixed-effects model revealed a significant main effect of time window (F(1,54) = 22.56, η²ₚ = 0.29, p < 0.001), indicating reduced alpha power immediately prior to target onset (Figure 4B). There was no main effect of group (F(1,55) = 0.135, p = 0.71) and no group × window interaction (F(1,54) = 1.65, p = 0.203), demonstrating comparable anticipatory alpha suppression in both groups when collapsed across conditions.

**Figure 4.**
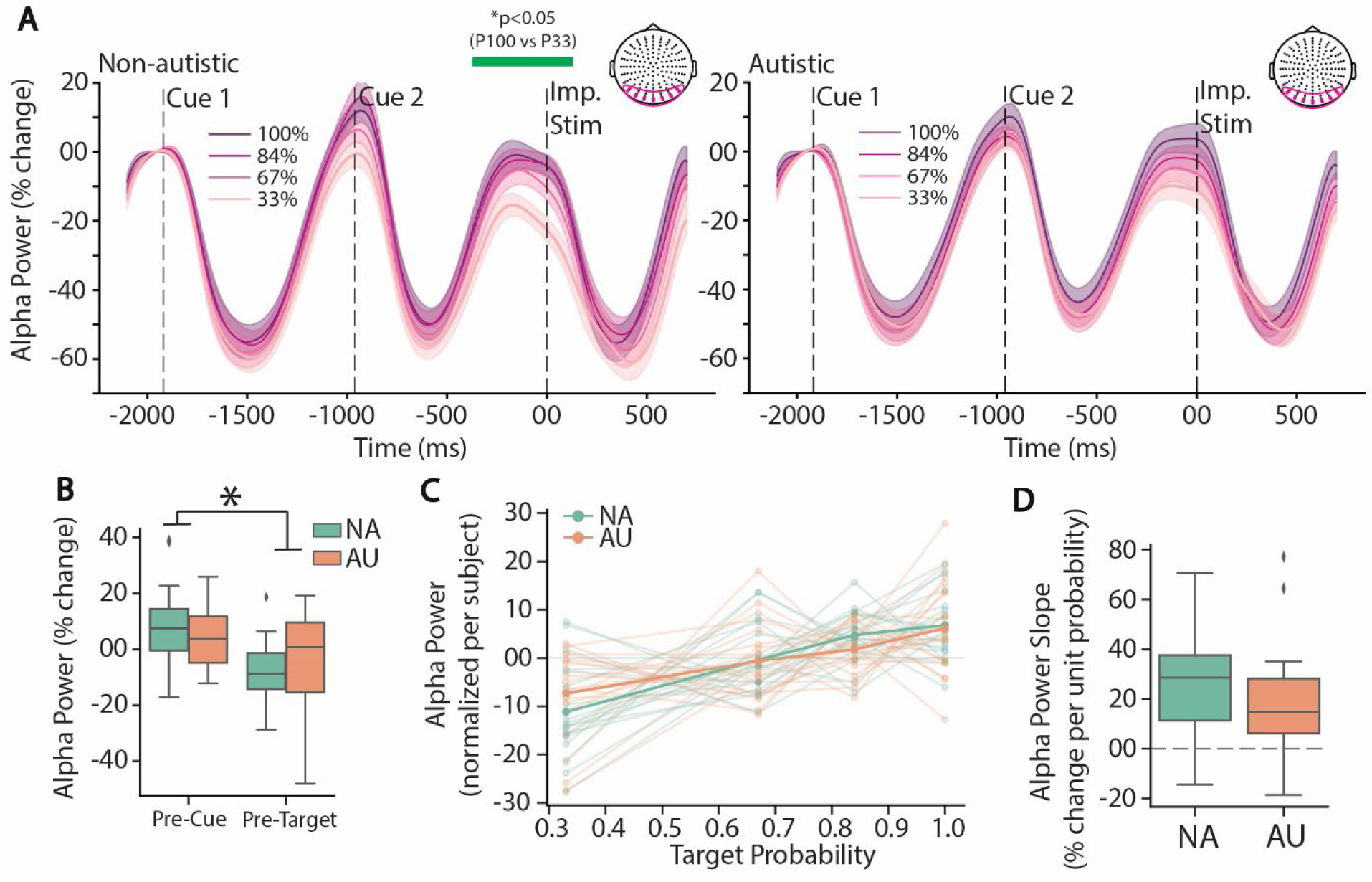
Similar modulation of induced alpha power in prediction of a low target probability in autistic and non-autistic groups. **(A)** Induced alpha power averaged over a cluster of parieto-occipital channels for non-autistic (left) and autistic group (right) for 100% (violet line), 84% (light violet), 67% (pink) and 33% target probability block (light pink). Green rectangle illustrates significant statistical differences assessed with cluster-based permutations (α = 0.05). (**B**) Average induced alpha power for the 100ms before Cue 2 (Pre-Cue) and for the 100ms before imperative stimulus onset (Pre-Target) over the cluster of central channels selected for the ERPs for non-autistic (green) and autistic (orange) averaged between the 4 targets probability conditions. * Indicates significant differences assessed by LMMs. **(C)** Linear regression of the induced alpha power slope by target probability with group-mean curve for non-autistic (blue) and autistic (orange) as individual regression curve in corresponding color. **(D)** Slope of the regression curve for each group.

We then assessed whether alpha modulation scaled with target probability. Temporal cluster-based permutation testing on the 33–100% difference wave revealed a significant alpha modulation in the non-autistic (NA) group, spanning −390ms to +170ms relative to target onset (Figure 4A). No significant clusters were observed in the autistic (AU) group. To quantify probability-dependent modulation, we extracted alpha power from the 100ms pre-target window for each condition and computed individual slopes across the four probability levels, paralleling the CNV analysis (Figure 4C). Slopes were significantly different from zero in both groups (NA: t = 5.61, p < 0.001; AU: t = 3.89, p < 0.001), indicating that alpha activity varied as a function of target probability. However, slope values did not significantly differ between groups (Figure 5C; t = 1.344, p = 0.187).

**Figure 5.**
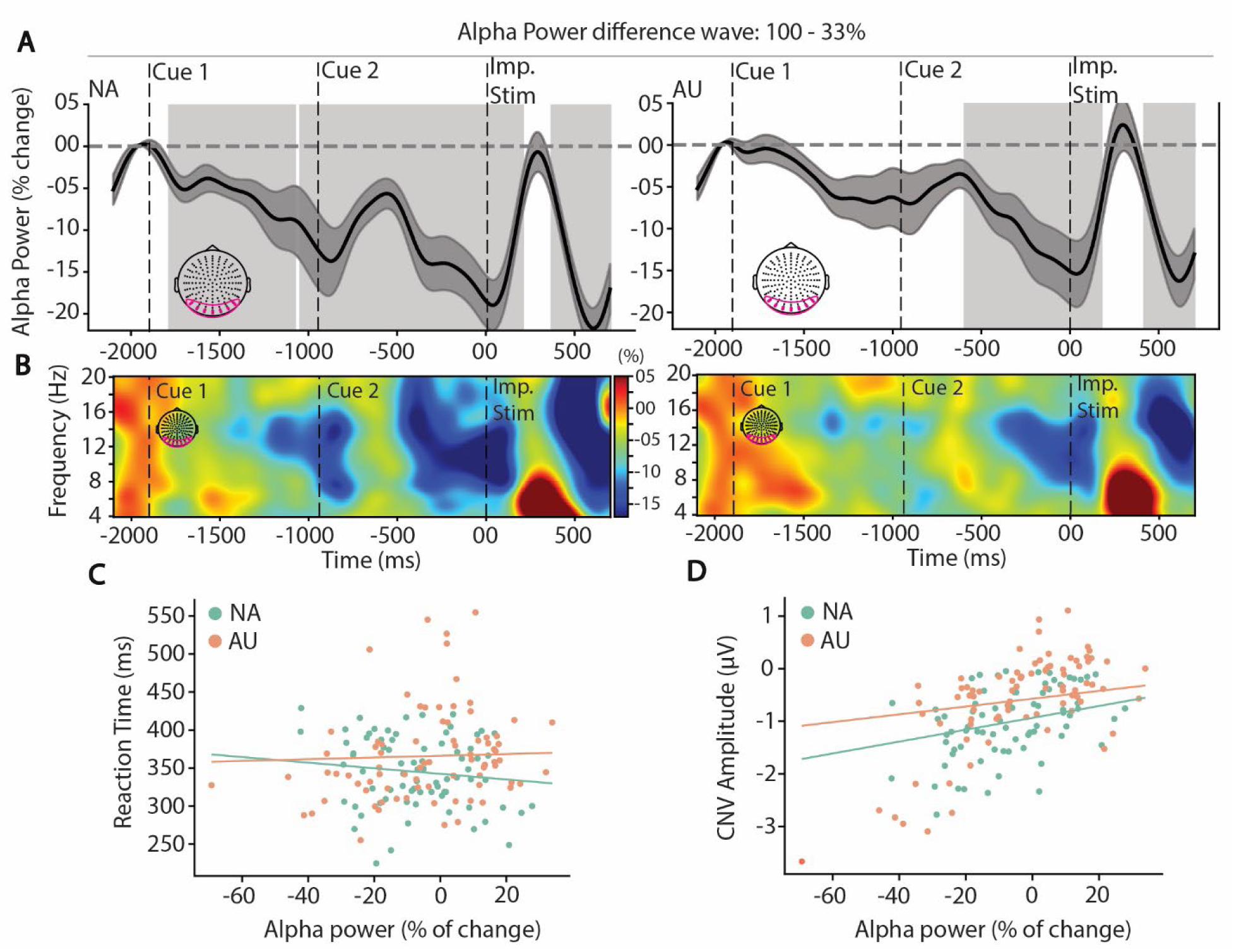
Temporal and spectral dynamic of anticipatory alpha modulation in the occipital cortex. **(A)** Time-course of the difference wave between 33% and 100% conditions showing the building-up of alpha desynchronization for non-autistic (left) and autistic (right) for a cluster of occipital channels and (**B**) corresponding time-frequency map of the difference wave. Dashed lines correspond to Cue 1 and Cue 2 and the imperative stimulus (‘Imp. Stim’; corresponding to both target and invalid stimulus) (**C**) Scatter plot between induced alpha power and Reaction time for non-autistic (blue) and autistic (red) and (**D**) between induced alpha power and CNV amplitude. Points represent condition-wise averages per participant. Solid lines depict linear mixed-effects model predictions (random intercept for Subject), controlling for target probability (Condition).

### Temporal Dynamics and Functional Relevance of Alpha Modulation

Because temporal cluster-based permutation testing indicated a stronger probability effect in the non-autistic (NA) group, we further examined the 33–100% alpha difference wave (Figure 5A). Significant clusters were observed in both groups; however, the onset of alpha modulation occurred substantially earlier in NA participants, following the onset of the first stimulus in the target sequence (beginning at −1790ms) than in autistic (AU) participants, where it was delayed until after the onset of the second stimulus in the sequence (at −560ms). This temporal shift suggests that although both groups exhibit anticipatory alpha modulation, its emergence is considerably delayed and temporally restricted in autism.

We next assessed whether induced alpha activity was functionally related to behavioral performance or to CNV amplitude. Linear mixed-effects models, controlling for target probability and repeated measures, revealed no significant association between alpha power (−100 to 0ms) and reaction time (β = 11.3ms per unit change in alpha, SE = 27.1, p = .677), and no significant Alpha × Group interaction (p = .168). Thus, there was no evidence that alpha modulation predicted behavioral performance in either group.

In contrast, alpha power was significantly associated with CNV amplitude (β = 0.74 µV, SE = 0.34, p = .029), such that greater alpha suppression (i.e., lower alpha power) predicted a more negative CNV. The absence of a significant Alpha × Group interaction (p = .359) indicates that this coupling between alpha desynchronization and CNV amplitude was comparable across autistic and non-autistic participants.

### Reduced P3b Modulation by Target Probability in Autism and Altered coupling with Anticipatory CNV

We next examined the P3b component elicited by target trials across all probability conditions. The P3b was extracted from a centro-parietal electrode cluster (Figure 6B; see Methods). To avoid baseline shifts related to the preceding CNV, we quantified P3b amplitude using a trough-to-peak measure (difference between the preceding negative deflection at 130–200ms and the P3b peak at 300–350ms).

**Figure 6.**
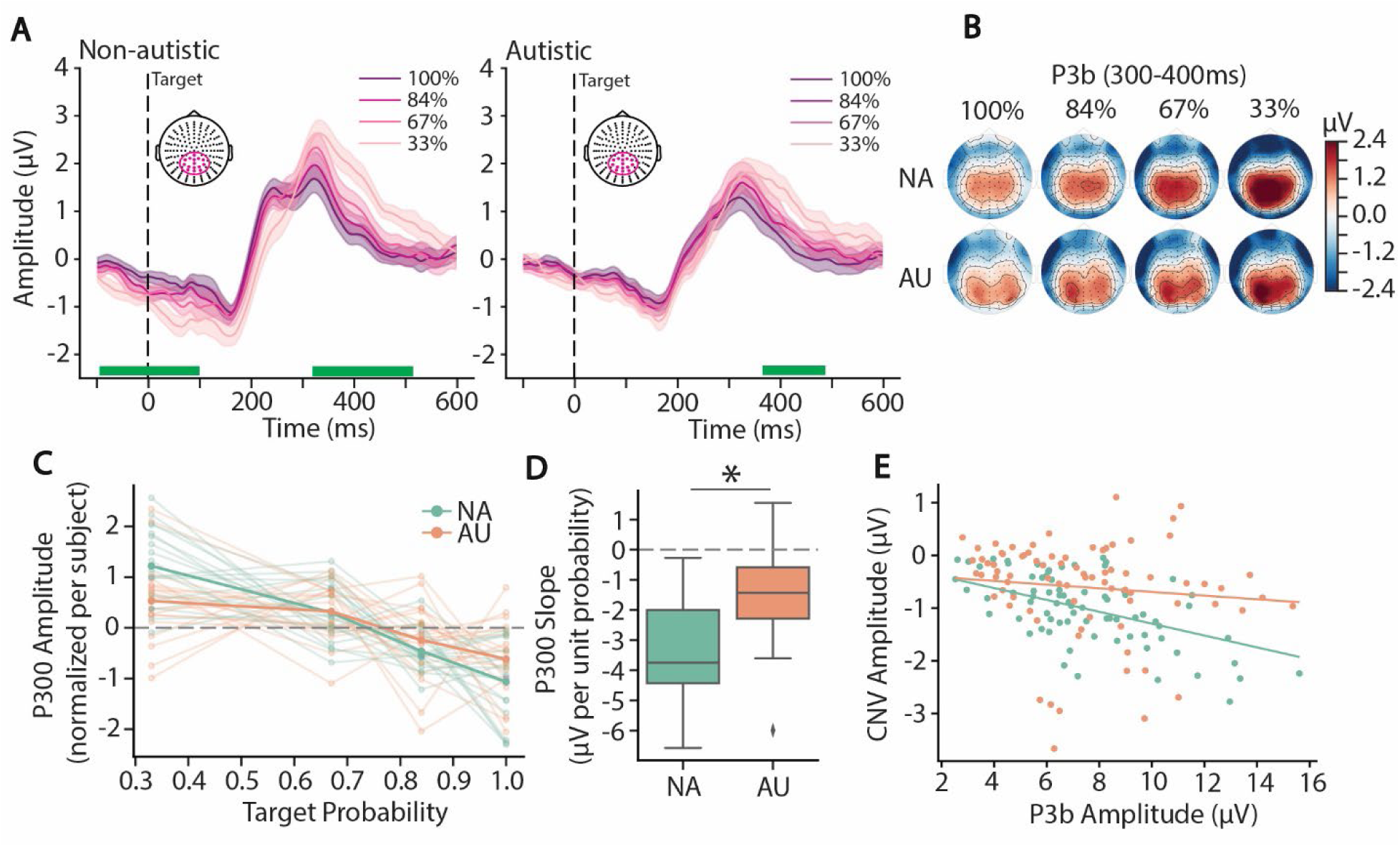
Reduced modulation of the P3b by target probability in autism. **(A)** Event-related potentials (ERPs) averaged over a cluster of centro-parietal channels in response to the target events for 100% (violet line), 84% (light violet), 67% (pink) and 33% target probability block (light pink), for non-autistic on the left and autistic on the right. Green rectangle illustrates significant statistical differences (significant only between 100 and 33%) assessed with cluster-based permutations (α = 0.05). **(B)** Topographical representation at the activity from 300-400ms after target onset from 100% on the left to 33% on the right, for non-autistic (top row) and autistic (bottom row). **(C)** Linear regression of the P300 slope by target probability with group-mean curve for non-autistic (blue) and autistic (orange) as individual regression curve in corresponding color. **(D)** Slope of the P300 regression curve for each group. * Indicates significant differences assessed by ANOVA. **(E)** Scatterplots showing the relationship between CNV and P300 amplitude for non-autistic (left) and autistic (right). Linear mixed model (LMM) regression lines are shown, with coefficients (β), standard errors (SE), and p-values reported from LMMs including subject as a random factor and target probability as a covariate.

Both groups exhibited a clear P3b across conditions (Figure 6A). A temporal cluster-based permutation test comparing the 100% and 33% probability conditions revealed significant differences in the non-autistic (NA) group both around target onset (−97 to +102ms), likely reflecting residual CNV activity, and during the P3b time window (321–498ms). In contrast, the autistic (AU) group showed significant differences only later in time (375–488ms). Topographical maps (300–400ms) demonstrated graded P3b modulation as a function of target probability, with larger amplitudes for lower-probability targets. Visually, this modulation appeared more pronounced in the NA group. To quantify contextual sensitivity, we computed individual slopes by regressing P3b amplitude onto target probability. Group comparison revealed significantly shallower slopes in the AU group (F(4, 36.9) = 7.97, p = 0.008), indicating reduced probability-dependent modulation. Nonetheless, slopes were significantly negative in both groups (NA: t = −7.95, p < 0.001; AU: t = −4.09, p < 0.001), demonstrating that P3b modulation by contextual probability was preserved but attenuated in autism.

Finally, we examined whether anticipatory activity predicted subsequent updating by modeling CNV amplitude as a function of P3b amplitude, controlling for probability condition and testing for group differences. The linear mixed-effects model revealed a significant P3b × Group interaction (β = −0.078, SE = 0.039, p = .047), indicating differential coupling between anticipatory preparation and target-related updating across groups. In NA participants, larger P3b amplitudes were associated with more negative CNV amplitudes (≈ −0.11 µV per unit P3b), consistent with functional coordination between anticipation and model updating. In contrast, this relationship was weak and non-significant in the AU group (β = −0.035, p = .277). These findings suggest that although anticipatory signals are present in both groups, their integration with subsequent target-related model updating processes is reduced in autism.

### Reduced P3a Amplitude and P3b Modulation to Invalid stimuli in Autism

We next examined neural responses to expectancy violations by focusing on invalid trials in which participants correctly withheld their response. These trials are conceptually similar to correct rejections in go/no-go paradigms, and our analyses therefore focused on the N2–P3a–P3b complex (see Methods), which is commonly examined in this context (Bokura et al., 2001; Polich, 2011).

The fronto-central N2 was clearly present in both groups, peaking around 270 ms (Figure 7A–D). Linear mixed-effects models (LMMs) conducted on participant-wise averages revealed no significant main effect of Condition or Group, and no interaction, indicating comparable N2 amplitudes across groups and probability contexts.

**Figure 7.**
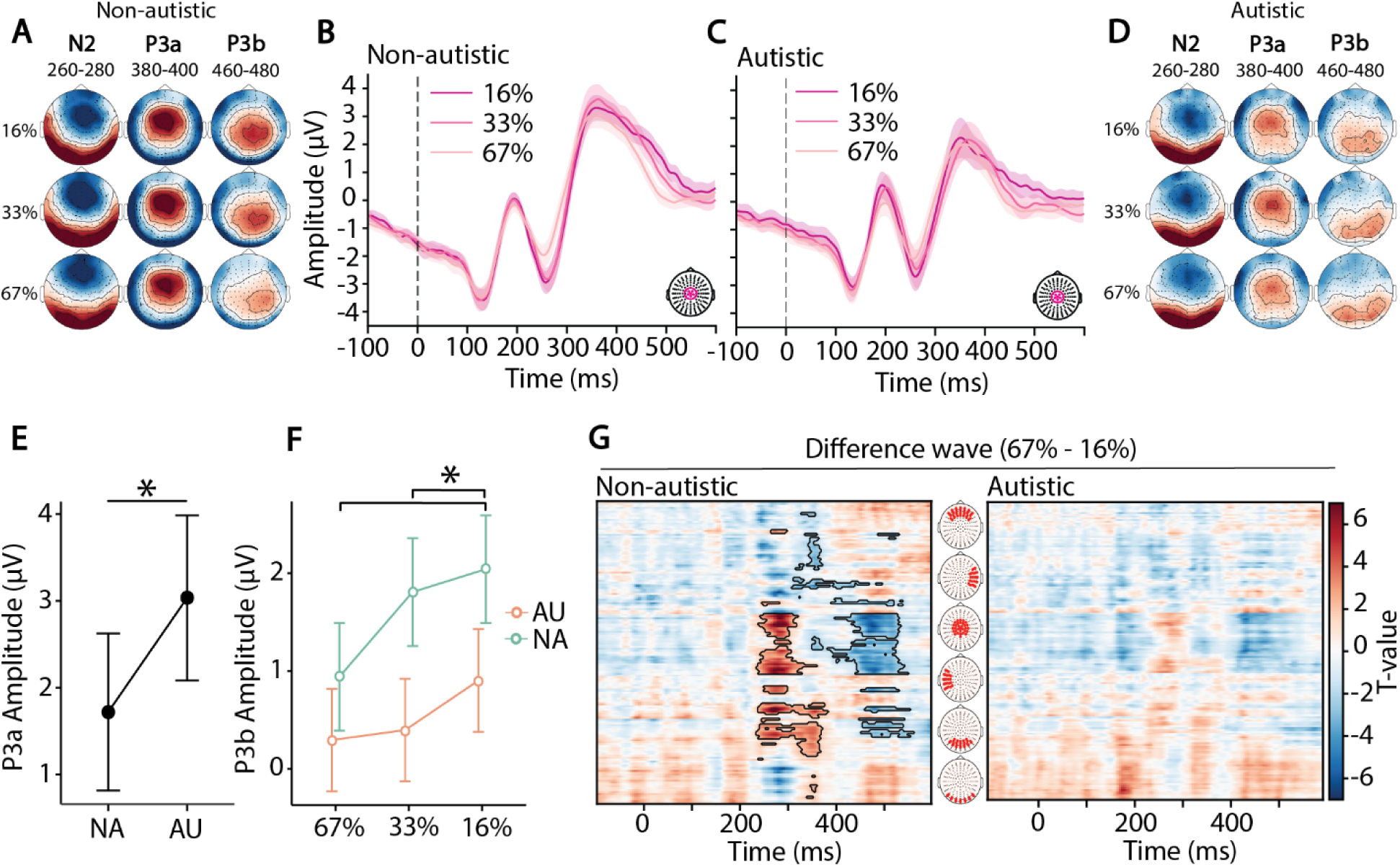
Reduced P3a amplitude and absence of P3b modulation to invalid stimuli in autism. **(A)** Topographical maps of neural responses to invalid stimuli for the N2 (260–280 ms), P3a (380–400 ms), and P3b (460–480 ms) across invalid-stimulus probabilities (16%: top row; 33%: middle row; 67%: bottom row) for the non-autistic group. **(D)** Corresponding topographical maps for the autistic group. **(B)** Event-related potentials (ERPs) averaged over the central electrode cluster for invalid trials in the non-autistic group, shown separately for 16% (light violet), 33% (pink), and 67% (light pink) invalid-stimulus probability blocks. **(C)** Corresponding ERPs for the autistic group. **(E)** Significant main effect of Group on mean P3a amplitude derived from linear mixed-effects modeling. **(F)** Significant Group × Condition interaction for P3b amplitude, reflecting reduced contextual modulation in the autistic group. **(G)** Spatio-temporal cluster-based permutation results for the 67%–16% difference wave in the non-autistic (left) and autistic (right) groups. Significant clusters are highlighted by a black outline.

The subsequent P3a, peaking around 390 ms over central electrodes (Figure 7A–D), showed a significant main effect of Group (Figure 7E; F(1,38) = 4.11, η²ₚ = 0.10, p = 0.050), with reduced amplitude in the autistic (AU) group relative to the non-autistic (NA) group. There was no significant main effect of Condition (F(2,76) = 2.58, p = 0.08) and no Group × Condition interaction (F(2,76) = 0.31, p = 0.73), suggesting that P3a amplitude did not scale with contextual certainty. The overall reduction in P3a amplitude in autism is consistent with an interpretation of diminished engagement of violation-sensitive or salience-related processes.

The later centro-parietal P3b, peaking around 470 ms (Figure 7A–D; see also Supplementary Fig. 1 for the ERP averaged over centro-parietal cluster, where P3b was greatest), revealed a significant main effect of Group (F(1,38) = 9.68, η²ₚ = 0.20, p = 0.004), reflecting reduced overall P3b amplitude in the AU group. A significant main effect of Condition was also observed (F(2,76) = 18.28, η²ₚ = 0.32, p < 0.001), along with a Group × Condition interaction (F(2,76) = 3.74, η²ₚ = 0.10, p = 0.02). Specifically, P3b amplitude increased as the probability of the invalid stimulus decreased in the NA group, whereas this contextual modulation was markedly attenuated in the AU group. Consistent with the reduced probability-dependent modulation observed for target-evoked P3b responses, these findings indicate diminished context-sensitive updating in autism when processing expectancy violations.

To complement these findings and avoid bias related to predefined electrodes or time windows, we conducted an exploratory spatio-temporal cluster-based permutation analysis comparing the two most extreme invalid-stimulus probability conditions (16% vs. 67%) within each group (Figure 7G). This data-driven approach revealed a significant spatio-temporal cluster in the non-autistic group only, encompassing central electrodes during the N2 and P3b time windows. No significant clusters were observed in the autistic group. These results confirm that contextual modulation of invalid-stimulus processing was present in the non-autistic group but markedly reduced in the autistic group.

## Discussion

Autistic individuals often resist even minor changes to everyday routines. One possibility is that this reflects difficulty flexibly adjusting internal predictive models in response to changing environmental contingencies. Within predictive processing frameworks, adaptive behavior depends not only on generating predictions but also on calibrating their certainty and updating them as contextual reliability shifts (Nobre et al., 2007; Rao & Ballard, 1999). We therefore tested the integrity of this adaptive system in autism by examining how different levels of predictive certainty modulate anticipatory activity and neural responses to both target and invalid events under graded uncertainty. We first investigated anticipatory mechanisms using two neural markers: the contingent negative variation (CNV) and occipital alpha-band desynchronization (α-ERD). We then examined model updating through the P3b to target and invalid stimuli, and prediction error–related processes via the N2/P3a complex to invalid stimuli. Four principal findings emerged. (1) Both autistic and non-autistic participants engaged in anticipatory mechanisms: CNV and α-ERD were robustly present prior to target onset. Greater alpha suppression predicted more negative CNV amplitudes, and more negative CNV amplitudes predicted faster reaction times, indicating that anticipatory signals were functionally relevant in both groups. (2) In response to targets, P3b amplitude scaled with target probability in both groups, being larger for less probable targets, consistent with probability-dependent updating. (3) However, autistic participants showed significantly reduced contextual modulation of anticipatory signals (CNV and alpha) and of the P3b. This attenuation was evident at the group and trial-level for the CNV, and for the P3b was particularly marked in response to invalid stimuli, where contextual modulation was largely absent in autism. Additionally, P3a amplitude to invalid stimuli was reduced overall in the autistic group. (4) Critically, the functional coupling between anticipatory preparation (CNV) and subsequent model updating (target-evoked P3b), observed in non-autistic participants, was disrupted in the autistic group. Together, these findings indicate that while core anticipatory mechanisms are present in autism, their tuning to contextual probabilities is less flexible, and their coordination with downstream updating processes is weakened. This pattern is consistent with reduced adaptive calibration of predictive processes.

### Anticipatory Mechanisms and Functional Relevance Across Groups

A central finding of the present study is that anticipatory neural mechanisms were clearly evident in both non-autistic and autistic participants, consistent with prior work (Sapey-Triomphe et al., 2023; Thillay et al., 2016). Both the contingent negative variation (CNV) and occipital alpha-band desynchronization (α-ERD) were robustly expressed across conditions, indicating intact engagement of preparatory neural processes in both groups. Critically, these anticipatory signals were functionally meaningful. CNV amplitude was significantly associated with reaction time in both groups, with more negative CNV amplitudes predicting faster responses. This relationship underscores the role of the CNV in motor and cognitive preparation that facilitates efficient behavioral responding (Kononowicz & Penney, 2016). The presence of this brain–behavior coupling in the autistic group demonstrates that anticipatory activity can effectively support performance when engaged, even though its contextual modulation was reduced relative to the non-autistic group.

Alpha ERD, commonly interpreted as reflecting increased cortical excitability and attentional allocation in sensory regions (Foxe & Snyder, 2011; Hanslmayr et al., 2011; Klimesch et al., 2007), was significantly associated with CNV amplitude, such that greater alpha suppression predicted a more negative CNV. However, alpha power did not directly predict reaction times in either group. This dissociation suggests that alpha ERD and the CNV index partially distinct yet coordinated stages of preparation. One plausible interpretation is that alpha desynchronization reflects modulation of sensory readiness, whereas the CNV captures motor and decision-related preparatory processes more directly tied to behavioral output. Taken together, these findings indicate that core anticipatory mechanisms are preserved in autism. The key difference observed in the present study lies not in the presence of preparatory processes per se, but in their flexibility and contextual tuning, which we address in the following sections.

### Reduced Adaptive Tuning of Anticipation and Updating in Autism

While anticipatory processes (alpha ERD and CNV) were present in the autistic group, autistic participants showed a markedly different pattern in how these processes were modulated by contextual probability and in how they interacted to support adaptive updating, as indexed by the P3. Across anticipatory, updating, and integrative levels, the autistic group demonstrated reduced flexibility in adjusting neural signals to changing environmental statistics. Thus, autistic participants had difficulty in dynamically tuning preparatory activity, weighting prediction errors, and coordinating these stages. This profile is consistent with altered predictive processing as a component of the autism phenotype.

### Attenuated Contextual Modulation of Anticipatory Processes (CNV, Alpha)

In the non-autistic control group, both CNV amplitude and alpha ERD scaled flexibly with contextual target probability, showing stronger anticipatory preparation under conditions requiring greater proactive control, consistent with previous studies linking these neural processes with flexible tuning of prediction certainty (Bagheri et al., 2025; Bauer et al., 1992; Bidet-Caulet et al., 2012; Gaillard, 1977; Korunka et al., 1993). In autism, this contextual modulation was significantly attenuated. Spatio-temporal cluster-based permutation testing further established that differences between the extreme conditions (100% vs. 33%) were limited in autism. Thus, although anticipatory processes were present, they were less dynamically shaped by changing environmental statistics in autism. These results could be considered to align with the hypo-prior hypothesis of predictive processing of autism, which suggests that autistic individuals rely less on top-down priors, leading to reduced contextual modulation and a greater reliance on incoming sensory input (Coll et al., 2020; Friston & Kiebel, 2009; Lawson et al., 2017; Lawson et al., 2014; Pellicano & Burr, 2012). However, they also suggest that on-line learning and model updating under changing circumstances are particularly altered. This interpretation resonates with behavioral findings from our study and others, showing smaller differences in RTs between high- and low-probability contexts in autism (Arthur et al., 2021; Perrykkad et al., 2021) but also neuroimaging studies showing reduced statistical learning using ERPs ((Jeste et al., 2015); and see (Bell et al., 2025) for a review of findings on statistical learning in autism). It also aligns with findings of slowed updating of internal representation in autism (Lieder et al., 2019; Vishne et al., 2021).

### Attenuated Contextual Modulation of Model (P3)

Probability-related modulation of the P3 also differed between the groups, for both target and invalid stimulus conditions. In the non-autistic control group, P3 amplitude to target events varied as a function of target probability in a manner consistent with its proposed role in model updating and prediction-error signaling (Bidet-Caulet et al., 2012; Donchin & Coles, 2010; Polich, 2011), with larger responses elicited by less probable events. In contrast, probability modulation of the P3 was greatly attenuated in autistic participants. A similar pattern was observed for the P300 response to an invalid stimulus (for invalid trials, the event that occurred in place of a target stimulus), with larger responses to invalid stimuli under high target probability contexts for non-autistic controls and notable reduction of this scaling in autism. This suggests diminished adjustment of internal models in autism when expectations are violated (i.e., when expectation differs from reality). This is consistent with altered precision-weighting of priors, which would hinder adaptive updating (Ishikawa et al., 2017; Lawson et al., 2017). Such a reduction in flexible contextual tuning provides a plausible mechanistic bridge to clinical features like resistance to change and intolerance of uncertainty, insofar as the system underweights contextual regularities and then fails to adjust internal models accordingly. Proper adjustment of the model requires implicitly calculating a running average of probabilities over time based on which updates are made.

### Altered Coordination Between Anticipation and Updating (CNV to P300)

In addition to revealing reduced modulation of anticipation and updating, the present findings also provide novel insight into how anticipatory mechanisms *interact* with model updating. In the non-autistic group, alpha ERD predicted CNV amplitude, and CNV amplitude predicted P300 amplitude, forming the appearance of a coordinated preparation-to-updating cascade. In autism, however, this chain of coordination appears disrupted. Although alpha ERD still predicted CNV amplitude, CNV no longer predicted P300 amplitude, indicating a decoupling between preparatory processes and subsequent cognitive updating. This breakdown may explain why anticipatory mechanisms, though present, fail to translate into flexible adaptation (Vishne et al., 2021). Without the ability to modulate the strength of the P300 “update” signal, internal models may remain overly rigid, consistent with predictive processing accounts of autism. This pattern highlights the importance of studying how neural processes interact in addition to considering isolated components and suggests that interventions aimed at improving prediction updating may need to target integration across neural systems (e.g., linking motor preparation with evaluative updating).

### Probability effects on the CNV

We found stronger anticipatory signals (more negative CNV and greater alpha ERD) in low-probability blocks, whereas some prior studies report larger anticipatory activity for high-probability cues for CNV (Bidet-Caulet et al., 2012; Feldman & Friston, 2010; Sznabel et al., 2023) and alpha power (Tarasi et al., 2022). We argue this discrepancy reflects differences in experimental design; our manipulation of cue-target probability was implicit and at the block level rather than that typically used when measuring probability effects on the CNV, in which cue-target probability varies is at the trial level, and is communicated by cue type. In the latter case, the cue on a given trial reliably signals the probability of an imminent response, promoting selective motor preparation, hence a larger CNV for high-probability cues. By contrast, our manipulation is contextual and held at the *block* level: the cue is constant and the likelihood of needing to respond varies across blocks. Under these conditions, low-probability blocks impose greater uncertainty and monitoring demands, encouraging sustained proactive control across the block, yielding a larger CNV (and stronger alpha suppression) as probability decreases. In short, we suggest that when validity is trial-wise, CNV scales with cue-locked motor set; when validity is contextual, CNV scales with tonic control under uncertainty. This pattern underscores the CNV’s multidimensional nature: it integrates certainty, task demands, and the balance between sustained control and selective, cue-locked preparation. Definitive confirmation of this explanation will require within-subject studies that directly compare trial-wise cue validity and block-level uncertainty in the same participants and such a design should be tested in future studies.

## Conclusion and Broader Implications

In summary, our findings demonstrate that autistic individuals engage top-down driven anticipatory mechanisms (CNV and alpha ERD) to optimize performance and that these top-down mechanisms retain their functional relevance for behavioral preparation. However, these mechanisms are less sensitive to contextual probability and do not drive effective model updating as indexed by the P300. This dual profile, robust preparation but reduced flexibility and integration, may underline key clinical features of autism, including insistence on sameness and difficulty adapting to change. From a theoretical perspective, these findings fit within predictive processing frameworks that emphasize the importance of modulating prediction precision based on environmental statistics (Cannon et al., 2021; Friston & Kiebel, 2009; Hesselmann et al., 2010). Future research should examine whether this inflexibility extends across modalities and contexts, and whether it is amenable to targeted interventions. Furthermore, future work should test whether these findings generalize to broader cohorts including individuals with lower cognitive or language abilities. Longitudinal studies could also determine whether disrupted coordination between model updating and anticipatory mechanisms contributes to the developmental trajectory of autism. By delineating how these processes are orchestrated, we move closer to understanding the neural mechanisms underlying adaptive flexibility, and its disruption in autism.

## Author contributions

S.R and S.M conceived the study. S.R and M.J.C implemented the experiment and recorded the data. T.V preprocessed the data and analyzed the data under the supervision of S.R. T.V and C.B wrote the first draft of the manuscript under the supervision of S.M.. M.J.C and S.M edited the manuscript.

## Acknowledgment

We are grateful to the individuals who participated in this research and their families for their time and their commitment to the advancement of scientific discovery; without them this work would not be possible. We would like to thank Drs. Catherine Sancimino and Juliana Bates, who administered or supervised the clinical assessments, and Drs. Jose Luis Pena, Ruben Coen Cagli, and Eric Hollander for their valuable inputs at S.R.’s student advisory committee meetings. This manuscript is based, in part, on work conducted in partial fulfillment of the requirements for the Ph.D. degree of Seydanir Reisli at the Albert Einstein College of Medicine.

The Human Clinical Phenotyping Core, where the majority of the children enrolled in this study were clinically evaluated, is a facility of the Rose F. Kennedy Intellectual and Developmental Disabilities Research Center (IDDRC) which is funded through a center grant from the Eunice Kennedy Shriver National Institute of Child Health & Human Development (NICHD U54 HD090260; P50 HD105352).

---

**Supplementary figure 1.**
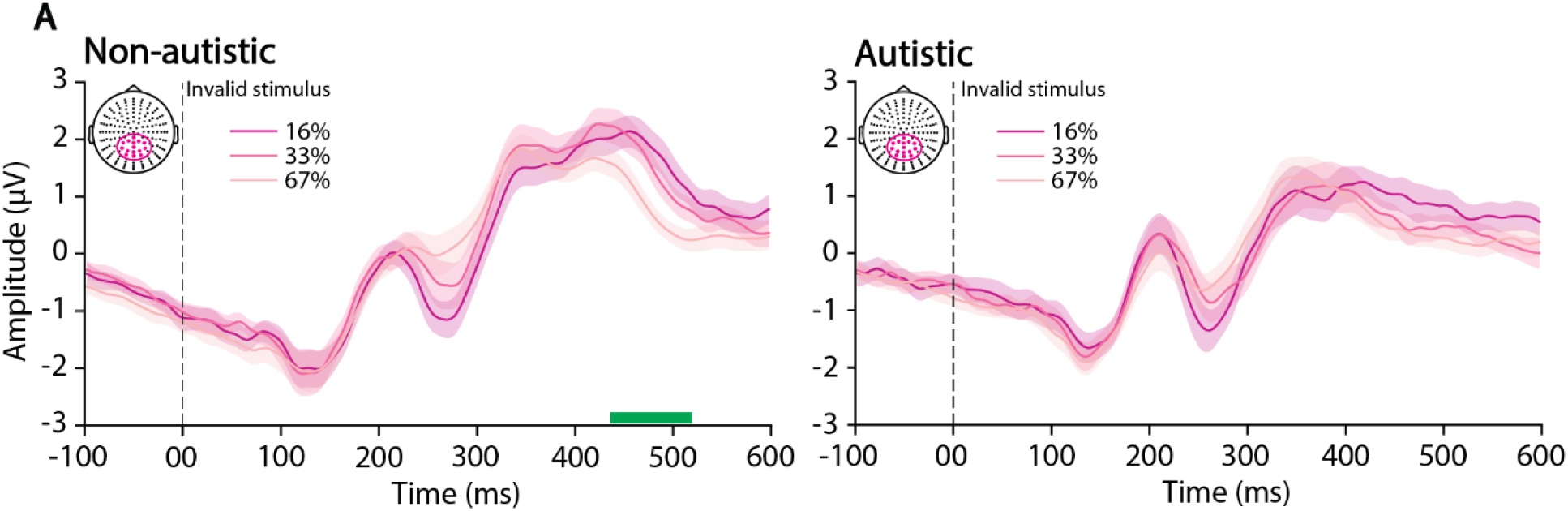
Reduced modulation of the P3b by invalid stimulus probability in autism. **(A)** Event-related potentials (ERPs) averaged over a cluster of centro-parietal channels in response to the invalid events for 16% (violet line), 33% (light violet), 67% (pink) for non-autistic on the left and autistic on the right. Green rectangle illustrates significant statistical differences (significant only between 16 and 67%) assessed with cluster-based permutations (α = 0.05).

**Supplementary table 1.**
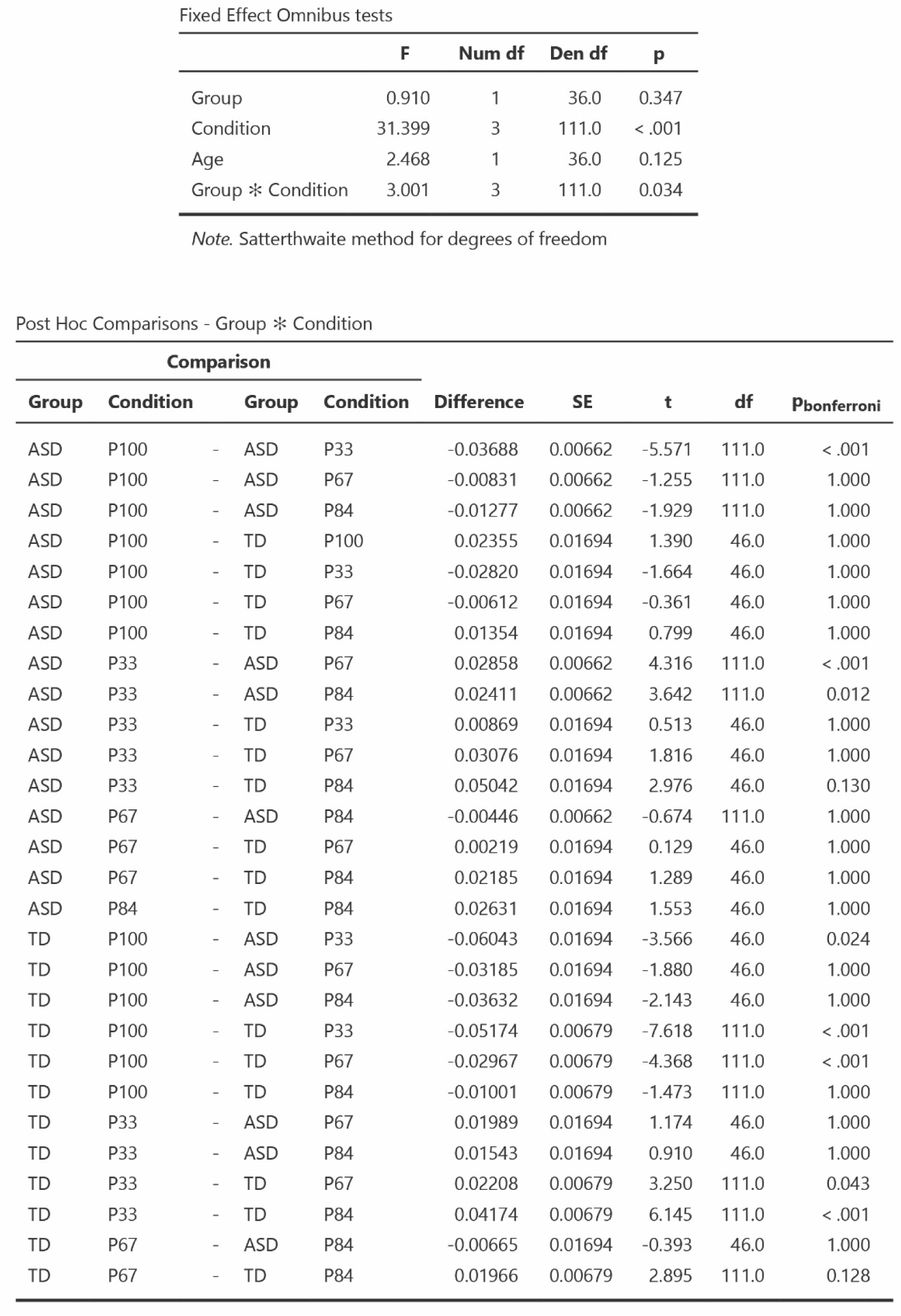
LMMs and post-hoc for reaction times results.

Fixed Effect Omnibus tests
|  | F | Num df | Den df | p |
| --- | --- | --- | --- | --- |
| Group | 0.910 | 1 | 36.0 | 0.347 |
| Condition | 31.399 | 3 | 111.0 | < .001 |
| Age | 2.468 | 1 | 36.0 | 0.125 |
| Group * Condition | 3.001 | 3 | 111.0 | 0.034 |

Note. Satterthwaite method for degrees of freedom

**Supplementary table 1. LMMs and post-hoc for reaction times results.**
| Comparison |  |  |  |  |  |  |  |  |
| --- | --- | --- | --- | --- | --- | --- | --- | --- |
| Group | Condition | Group | Condition | Difference | SE | t | df | Pbonferroni |
| ASD | P100 | - ASD | P33 | -0.03688 | 0.00662 | -5.571 | 111.0 | < .001 |
| ASD | P100 | - ASD | P67 | -0.00831 | 0.00662 | -1.255 | 111.0 | 1.000 |
| ASD | P100 | - ASD | P84 | -0.01277 | 0.00662 | -1.929 | 111.0 | 1.000 |
| ASD | P100 | - TD | P100 | 0.02355 | 0.01694 | 1.390 | 46.0 | 1.000 |
| ASD | P100 | - TD | P33 | -0.02820 | 0.01694 | -1.664 | 46.0 | 1.000 |
| ASD | P100 | - TD | P67 | -0.00612 | 0.01694 | -0.361 | 46.0 | 1.000 |
| ASD | P100 | - TD | P84 | 0.01354 | 0.01694 | 0.799 | 46.0 | 1.000 |
| ASD | P33 | - ASD | P67 | 0.02858 | 0.00662 | 4.316 | 111.0 | < .001 |
| ASD | P33 | - ASD | P84 | 0.02411 | 0.00662 | 3.642 | 111.0 | 0.012 |
| ASD | P33 | - TD | P33 | 0.00869 | 0.01694 | 0.513 | 46.0 | 1.000 |
| ASD | P33 | - TD | P67 | 0.03076 | 0.01694 | 1.816 | 46.0 | 1.000 |
| ASD | P33 | - TD | P84 | 0.05042 | 0.01694 | 2.976 | 46.0 | 0.130 |
| ASD | P67 | - ASD | P84 | -0.00446 | 0.00662 | -0.674 | 111.0 | 1.000 |
| ASD | P67 | - TD | P67 | 0.00219 | 0.01694 | 0.129 | 46.0 | 1.000 |
| ASD | P67 | - TD | P84 | 0.02185 | 0.01694 | 1.289 | 46.0 | 1.000 |
| ASD | P84 | - TD | P84 | 0.02631 | 0.01694 | 1.553 | 46.0 | 1.000 |
| TD | P100 | - ASD | P33 | -0.06043 | 0.01694 | -3.566 | 46.0 | 0.024 |
| TD | P100 | - ASD | P67 | -0.03185 | 0.01694 | -1.880 | 46.0 | 1.000 |
| TD | P100 | - ASD | P84 | -0.03632 | 0.01694 | -2.143 | 46.0 | 1.000 |
| TD | P100 | - TD | P33 | -0.05174 | 0.00679 | -7.618 | 111.0 | < .001 |
| TD | P100 | - TD | P67 | -0.02967 | 0.00679 | -4.368 | 111.0 | < .001 |
| TD | P100 | - TD | P84 | -0.01001 | 0.00679 | -1.473 | 111.0 | 1.000 |
| TD | P33 | - ASD | P67 | 0.01989 | 0.01694 | 1.174 | 46.0 | 1.000 |
| TD | P33 | - ASD | P84 | 0.01543 | 0.01694 | 0.910 | 46.0 | 1.000 |
| TD | P33 | - TD | P67 | 0.02208 | 0.00679 | 3.250 | 111.0 | 0.043 |
| TD | P33 | - TD | P84 | 0.04174 | 0.00679 | 6.145 | 111.0 | < .001 |
| TD | P67 | - ASD | P84 | -0.00665 | 0.01694 | -0.393 | 46.0 | 1.000 |
| TD | P67 | - TD | P84 | 0.01966 | 0.00679 | 2.895 | 111.0 | 0.128 |

## References

American Psychiatric, A. (2013). DIAGNOSTIC AND STATISTICAL MANUAL OF MENTAL DISORDERS (5 ed.).

Arthur, T., Harris, D., Buckingham, G., Brosnan, M., Wilson, M., Williams, G., & Vine, S. (2021). An examination of active inference in autistic adults using immersive virtual reality. Sci Rep, 11(1), 20377. 10.1038/s41598-021-99864-y

Bagheri, S., Cardinaux, A., Bungert, L., Li, C., O’Brien, A., Sinha, P., & Cannon, J. (2025). Reduced anticipatory motor preparation and altered EEG signatures of prediction in autistic adults. 10.31234/osf.io/297tq_v1

Bar, M. (2007). The proactive brain: using analogies and associations to generate predictions. Trends Cogn Sci, 11(7), 280–289. 10.1016/j.tics.2007.05.005

Bar, M., Kassam, K. S., Ghuman, A. S., Boshyan, J., Schmid, A. M., Dale, A. M., Hamalainen, M. S., Marinkovic, K., Schacter, D. L., Rosen, B. R., & Halgren, E. (2006). Top-down facilitation of visual recognition. Proc Natl Acad Sci U S A, 103(2), 449–454. 10.1073/pnas.0507062103

Bauer, H., Rebert, C., Korunka, C., & Leodolter, M. (1992). Rare events and the CNV--the oddball CNV. Int J Psychophysiol, 13(1), 51–58. 10.1016/0167-8760(92)90020-c

Beker, S., Foxe, J. J., & Molholm, S. (2021). Oscillatory entrainment mechanisms and anticipatory predictive processes in children with autism spectrum disorder. J Neurophysiol, 126(5), 1783–1798. 10.1152/jn.00329.2021

Bell, R. R., Thomas, H. R., Saffran, J. R., & Eigsti, I. M. (2025). A systematic review of statistical learning in autism spectrum disorder. Mol Autism, 17(1), 2. 10.1186/s13229-025-00697-7

Bidet-Caulet, A., Barbe, P. G., Roux, S., Viswanath, H., Barthelemy, C., Bruneau, N., Knight, R. T., & Bonnet-Brilhault, F. (2012). Dynamics of anticipatory mechanisms during predictive context processing. Eur J Neurosci, 36(7), 2996–3004. 10.1111/j.1460-9568.2012.08223.x

Bigdely-Shamlo, N., Mullen, T., Kothe, C., Su, K. M., & Robbins, K. A. (2015). The PREP pipeline: Standardized preprocessing for large-scale EEG analysis. Frontiers in Neuroinformatics, 9(JUNE), 1–19. 10.3389/fninf.2015.00016

Boettcher, S. E. P., Stokes, M. G., Nobre, A. C., & van Ede, F. (2020). One Thing Leads to Another: Anticipating Visual Object Identity Based on Associative-Memory Templates. J Neurosci, 40(20), 4010–4020. 10.1523/JNEUROSCI.2751-19.2020

Bokura, H., Yamaguchi, S., & Kobayashi, S. (2001). Electrophysiological correlates for response inhibition in a Go/NoGo task. Clin Neurophysiol, 112(12), 2224–2232. 10.1016/s1388-2457(01)00691-5

Cannon, J., O’Brien, A. M., Bungert, L., & Sinha, P. (2021). Prediction in Autism Spectrum Disorder: A Systematic Review of Empirical Evidence. Autism Res, 14(4), 604–630. 10.1002/aur.2482

Chambon, V., Farrer, C., Pacherie, E., Jacquet, P. O., Leboyer, M., & Zalla, T. (2017). Reduced sensitivity to social priors during action prediction in adults with autism spectrum disorders. Cognition, 160, 17–26. 10.1016/j.cognition.2016.12.005

Chen, T. C., Hsieh, M. H., Lin, Y. T., Chan, P. S., & Cheng, C. H. (2020). Mismatch negativity to different deviant changes in autism spectrum disorders: A meta-analysis. Clin Neurophysiol, 131(3), 766–777. 10.1016/j.clinph.2019.10.031

Cohen, M. X. (2014). Fluctuations in oscillation frequency control spike timing and coordinate neural networks. Journal of Neuroscience, 34(27), 8988–8998. 10.1523/JNEUROSCI.0261-14.2014

Coll, M. P., Whelan, E., Catmur, C., & Bird, G. (2020). Autistic traits are associated with atypical precision-weighted integration of top-down and bottom-up neural signals. Cognition, 199, 104236. 10.1016/j.cognition.2020.104236

Donchin, E., & Coles, M. G. H. (2010). Is the P300 component a manifestation of context updating? Behavioral and Brain Sciences, 11(3), 357–374. 10.1017/s0140525x00058027

Feldman, H., & Friston, K. J. (2010). Attention, uncertainty, and free-energy. Front Hum Neurosci, 4, 215. 10.3389/fnhum.2010.00215

Folstein, J. R., & Van Petten, C. (2008). Influence of cognitive control and mismatch on the N2 component of the ERP: a review. Psychophysiology, 45(1), 152–170. 10.1111/j.1469-8986.2007.00602.x

Foxe, J. J., & Snyder, A. C. (2011). The role of alpha-band brain oscillations as a sensory suppression mechanism during selective attention. Frontiers in Psychology, 2(JUL). 10.3389/fpsyg.2011.00154

Friston, K., & Kiebel, S. (2009). Predictive coding under the free-energy principle. Philos Trans R Soc Lond B Biol Sci, 364(1521), 1211–1221. 10.1098/rstb.2008.0300

Gaillard, A. W. (1977). The late CNV wave: preparation versus expectancy. Psychophysiology, 14(6), 563–568. 10.1111/j.1469-8986.1977.tb01200.x

Galvao-Carmona, A., Gonzalez-Rosa, J. J., Hidalgo-Munoz, A. R., Paramo, D., Benitez, M. L., Izquierdo, G., & Vazquez-Marrufo, M. (2014). Disentangling the attention network test: behavioral, event related potentials, and neural source analyses. Front Hum Neurosci, 8, 813. 10.3389/fnhum.2014.00813

Garrido, M. I., Kilner, J. M., Stephan, K. E., & Friston, K. J. (2009). The mismatch negativity: a review of underlying mechanisms. Clin Neurophysiol, 120(3), 453–463. 10.1016/j.clinph.2008.11.029

Gomot, M., & Wicker, B. (2012). A challenging, unpredictable world for people with autism spectrum disorder. Int J Psychophysiol, 83(2), 240–247. 10.1016/j.ijpsycho.2011.09.017

Gould, I. C., Rushworth, M. F., & Nobre, A. C. (2011). Indexing the graded allocation of visuospatial attention using anticipatory alpha oscillations. J Neurophysiol, 105(3), 1318–1326. 10.1152/jn.00653.2010

Gramfort, A., Luessi, M., Larson, E., Engemann, D. A., Strohmeier, D., Brodbeck, C., Goj, R., Jas, M., Brooks, T., Parkkonen, L., & Hamalainen, M. (2013). MEG and EEG data analysis with MNE-Python. Front Neurosci, 7, 267. 10.3389/fnins.2013.00267

Gregory, R. L. (1980). Perceptions as hypotheses. Philos Trans R Soc Lond B Biol Sci, 290(1038), 181–197. 10.1098/rstb.1980.0090

Groppe, D. M., Urbach, T. P., & Kutas, M. (2011). Mass univariate analysis of event-related brain potentials/fields I: a critical tutorial review. Psychophysiology, 48(12), 1711–1725. 10.1111/j.1469-8986.2011.01273.x

Hanslmayr, S., Gross, J., Klimesch, W., & Shapiro, K. L. (2011). The role of alpha oscillations in temporal attention. Brain Res Rev, 67(1-2), 331–343. 10.1016/j.brainresrev.2011.04.002

He, B., Lian, J., Spencer, K. M., Dien, J., & Donchin, E. (2001). A cortical potential imaging analysis of the P300 and novelty P3 components. Hum Brain Mapp, 12(2), 120–130. 10.1002/1097-0193(200102)12:2<120::aid-hbm1009>3.0.co;2-v

Hesselmann, G., Sadaghiani, S., Friston, K. J., & Kleinschmidt, A. (2010). Predictive coding or evidence accumulation? False inference and neuronal fluctuations. PLoS ONE, 5(3), e9926. 10.1371/journal.pone.0009926

Hohwy, J. (2017). Priors in perception: Top-down modulation, Bayesian perceptual learning rate, and prediction error minimization. Conscious Cogn, 47, 75–85. 10.1016/j.concog.2016.09.004

Ishikawa, M., Itakura, S., & Tanabe, H. C. (2017). Autistic Traits Affect P300 Response to Unexpected Events, regardless of Mental State Inferences. Autism Res Treat, 2017, 8195129. 10.1155/2017/8195129

Jeste, S. S., Kirkham, N., Senturk, D., Hasenstab, K., Sugar, C., Kupelian, C., Baker, E., Sanders, A. J., Shimizu, C., Norona, A., Paparella, T., Freeman, S. F., & Johnson, S. P. (2015). Electrophysiological evidence of heterogeneity in visual statistical learning in young children with ASD. Dev Sci, 18(1), 90–105. 10.1111/desc.12188

Kalcher, J., & Pfurtscheller, G. (1995). Discrimination between phase-locked and non-phase-locked event-related EEG activity. Electroencephalogr Clin Neurophysiol, 94(5), 381–384. 10.1016/0013-4694(95)00040-6

Keller, G. B., & Mrsic-Flogel, T. D. (2018). Predictive Processing: A Canonical Cortical Computation. Neuron, 100(2), 424–435. 10.1016/j.neuron.2018.10.003

Klimesch, W., Sauseng, P., & Hanslmayr, S. (2007). EEG alpha oscillations: The inhibition-timing hypothesis. In Brain Research Reviews (Vol. 53, pp. 63-88).

Kononowicz, T. W., & Penney, T. B. (2016). The contingent negative variation (CNV): timing isn’t everything. Current Opinion in Behavioral Sciences, 8, 231–237. 10.1016/j.cobeha.2016.02.022

Korunka, C., Wenzel, T., & Bauer, H. (1993). The ‘oddball CNV’ as an indicator of different information processing in patients with panic disorders. Int J Psychophysiol, 15(3), 207–215. 10.1016/0167-8760(93)90004-9

Lasaponara, S., D’Onofrio, M., Pinto, M., Dragone, A., Menicagli, D., Bueti, D., De Lucia, M., Tomaiuolo, F., & Doricchi, F. (2018). EEG Correlates of Preparatory Orienting, Contextual Updating, and Inhibition of Sensory Processing in Left Spatial Neglect. J Neurosci, 38(15), 3792–3808. 10.1523/JNEUROSCI.2817-17.2018

Lawson, R. P., Mathys, C., & Rees, G. (2017). Adults with autism overestimate the volatility of the sensory environment. Nat Neurosci, 20(9), 1293–1299. 10.1038/nn.4615

Lawson, R. P., Rees, G., & Friston, K. J. (2014). An aberrant precision account of autism. Front Hum Neurosci, 8, 302. 10.3389/fnhum.2014.00302

Li, S., Seger, C. A., Zhang, J., Liu, M., Dong, W., Liu, W., & Chen, Q. (2023). Alpha oscillations encode Bayesian belief updating underlying attentional allocation in dynamic environments. NeuroImage, 284, 120464. 10.1016/j.neuroimage.2023.120464

Lieder, I., Adam, V., Frenkel, O., Jaffe-Dax, S., Sahani, M., & Ahissar, M. (2019). Perceptual bias reveals slow-updating in autism and fast-forgetting in dyslexia. Nat Neurosci, 22(2), 256–264. 10.1038/s41593-018-0308-9

Lord, C., Rutter, M., & Lecouteur, A. (1994). Autism Diagnostic Interview-Revised - a Revised Version of a Diagnostic Interview for Caregivers of Individuals with Possible Pervasive Developmental Disorders. Journal of Autism and Developmental Disorders, 24(5), 659–685. Doi 10.1007/Bf02172145

Maenner, M. J., Warren, Z., Williams, A. R., Amoakohene, E., Bakian, A. V., Bilder, D. A., Durkin, M. S., Fitzgerald, R. T., Furnier, S. M., Hughes, M. M., Ladd-Acosta, C. M., McArthur, D., Pas, E. T., Salinas, A., Vehorn, A., Williams, S., Esler, A., Grzybowski, A., Hall-Lande, J., … Shaw, K. A. (2023). Prevalence and Characteristics of Autism Spectrum Disorder Among Children Aged 8 Years - Autism and Developmental Disabilities Monitoring Network, 11 Sites, United States, 2020. MMWR Surveill Summ, 72(2), 1–14. 10.15585/mmwr.ss7202a1

Maris, E., & Oostenveld, R. (2007). Nonparametric statistical testing of EEG- and MEG-data. Journal of Neuroscience Methods, 164(1), 177–190. 10.1016/j.jneumeth.2007.03.024

Michelini, G., Kitsune, V., Vainieri, I., Hosang, G. M., Brandeis, D., Asherson, P., & Kuntsi, J. (2018). Shared and Disorder-Specific Event-Related Brain Oscillatory Markers of Attentional Dysfunction in ADHD and Bipolar Disorder. Brain Topogr, 31(4), 672–689. 10.1007/s10548-018-0625-z

Nobre, A., Correa, A., & Coull, J. (2007). The hazards of time. Curr Opin Neurobiol, 17(4), 465–470. 10.1016/j.conb.2007.07.006

Palmer, C. J., Seth, A. K., & Hohwy, J. (2015). The felt presence of other minds: Predictive processing, counterfactual predictions, and mentalising in autism. Conscious Cogn, 36, 376–389. 10.1016/j.concog.2015.04.007

Pellicano, E., & Burr, D. (2012). When the world becomes ‘too real’: a Bayesian explanation of autistic perception. Trends Cogn Sci, 16(10), 504–510. 10.1016/j.tics.2012.08.009

Perrin, Pernier, Bertrand, & Echallier. (1989). Spherical splines for scalp potential and current density mapping. Electroencephalography and Clinical Neurophysiology, 72, 184–187.

Perrykkad, K., Lawson, R. P., Jamadar, S., & Hohwy, J. (2021). The effect of uncertainty on prediction error in the action perception loop. Cognition, 210, 104598. 10.1016/j.cognition.2021.104598

Pesthy, O., Farkas, K., Sapey-Triomphe, L. A., Guttengeber, A., Komoroczy, E., Janacsek, K., Rethelyi, J. M., & Nemeth, D. (2023). Intact predictive processing in autistic adults: evidence from statistical learning. Sci Rep, 13(1), 11873. 10.1038/s41598-023-38708-3

Pfurtscheller, G., & Klimesch, W. (1992). Functional Topography During a Visuoverbal Judgment Task Studied with Event-Related Desynchronization Mapping. Journal of Clinical Neurophysiology, 9(1). 10.1097/00004691-199201000-00013

Polich, J. (2007). Updating P300: an integrative theory of P3a and P3b. Clin Neurophysiol, 118(10), 2128–2148. 10.1016/j.clinph.2007.04.019

Polich, J. (2011). Neuropsychology of P300. In The Oxford Handbook of Event-Related Potential Components (pp. 0). Oxford University Press. 10.1093/oxfordhb/9780195374148.013.0089

Rao, R. P., & Ballard, D. H. (1999). Predictive coding in the visual cortex: a functional interpretation of some extra-classical receptive-field effects. Nat Neurosci, 2(1), 79–87. 10.1038/4580

Rohenkohl, G., & Nobre, A. C. (2011). alpha oscillations related to anticipatory attention follow temporal expectations. J Neurosci, 31(40), 14076–14084. 10.1523/JNEUROSCI.3387-11.2011

Rong, Y., Yang, C.-J., Jin, Y., & Wang, Y. (2021). Prevalence of attention-deficit/hyperactivity disorder in individuals with autism spectrum disorder: A meta-analysis. Research in Autism Spectrum Disorders, 83. 10.1016/j.rasd.2021.101759

Sapey-Triomphe, L. A., Bouet, R., Mattout, J., Sonie, S., Schmitz, C., & Lecaignard, F. (2025). Systematic Review and Meta-Analysis of Mismatch Negativity in Autism: Insights Into Predictive Mechanisms. Autism Res, 18(12), 2431–2450. 10.1002/aur.70131

Sapey-Triomphe, L. A., Pattyn, L., Weilnhammer, V., Sterzer, P., & Wagemans, J. (2023). Neural correlates of hierarchical predictive processes in autistic adults. Nat Commun, 14(1), 3640. 10.1038/s41467-023-38580-9

Sapey-Triomphe, L. A., Timmermans, L., & Wagemans, J. (2021). Priors Bias Perceptual Decisions in Autism, But Are Less Flexibly Adjusted to the Context. Autism Res, 14(6), 1134–1146. 10.1002/aur.2452

Schuller, T., Mengotti, P., Zabicki, A., Huys, D., Barbe, M. T., Fink, G. R., Visser-Vandewalle, V., Vossel, S., & Baldermann, J. C. (2025). Alterations of sensorimotor predictive processes and their electrophysiological signatures in Tourette syndrome. Brain Commun, 7(6), fcaf458. 10.1093/braincomms/fcaf458

Schwartz, S., Shinn-Cunningham, B., & Tager-Flusberg, H. (2018). Meta-analysis and systematic review of the literature characterizing auditory mismatch negativity in individuals with autism. Neurosci Biobehav Rev, 87, 106–117. 10.1016/j.neubiorev.2018.01.008

Shi, Z., Allenmark, F., Theisinger, L. A., Pistorius, R. L., Glasauer, S., Muller, H. J., & Falter-Wagner, C. M. (2025). Predictive Processing in Autism Spectrum Disorder: The Atypical Iterative Prior Updating Account. Biol Psychiatry Glob Open Sci, 5(3), 100468. 10.1016/j.bpsgos.2025.100468

Simard, I., Luck, D., Mottron, L., Zeffiro, T. A., & Soulieres, I. (2015). Autistic fluid intelligence: Increased reliance on visual functional connectivity with diminished modulation of coupling by task difficulty. Neuroimage Clin, 9, 467–478. 10.1016/j.nicl.2015.09.007

Smith, M. E., Halgren, E., Sokolik, M., Baudena, P., Musolino, A., Liegeois-Chauvel, C., & Chauvel, P. (1990). The intracranial topography of the P3 event-related potential elicited during auditory oddball. Electroencephalogr Clin Neurophysiol, 76(3), 235–248. 10.1016/0013-4694(90)90018-f

Squires, N. K., Squires, K. C., & Hillyard, S. A. (1975). Two varieties of long-latency positive waves evoked by unpredictable auditory stimuli in man. Electroencephalogr Clin Neurophysiol, 38(4), 387–401. 10.1016/0013-4694(75)90263-1

Sznabel, D., Land, R., Kopp, B., & Kral, A. (2023). The relation between implicit statistical learning and proactivity as revealed by EEG. Sci Rep, 13(1), 15787. 10.1038/s41598-023-42116-y

Tarasi, L., di Pellegrino, G., & Romei, V. (2022). Are you an empiricist or a believer? Neural signatures of predictive strategies in humans. Prog Neurobiol, 219, 102367. 10.1016/j.pneurobio.2022.102367

Tecce, J. J. (1972). Contingent negative variation (CNV) and psychological processes in man. Psychological Bulletin, 77(2), 73–108. 10.1037/h0032177

Thillay, A., Lemaire, M., Roux, S., Houy-Durand, E., Barthelemy, C., Knight, R. T., Bidet-Caulet, A., & Bonnet-Brilhault, F. (2016). Atypical Brain Mechanisms of Prediction According to Uncertainty in Autism. Front Neurosci, 10, 317. 10.3389/fnins.2016.00317

Van de Cruys, S., Evers, K., Van der Hallen, R., Van Eylen, L., Boets, B., de-Wit, L., & Wagemans, J. (2014). Precise minds in uncertain worlds: Predictive coding in autism. Psychological Review, 121(4), 649–675. 10.1037/a0037665

Vishne, G., Jacoby, N., Malinovitch, T., Epstein, T., Frenkel, O., & Ahissar, M. (2021). Slow update of internal representations impedes synchronization in autism. Nat Commun, 12(1), 5439. 10.1038/s41467-021-25740-y

Walter, W. G., Cooper, R., Aldridge, V., McCallum, W., & Winter, A. (1964). Contingent negative variation: an electric sign of sensori-motor association and expectancy in the human brain. Nature, 203(4943), 380–384.

Wang, Q., She, S., Luo, L., Li, H., Ning, Y., Ren, J., Wu, Z., Huang, R., & Zheng, Y. (2020). Abnormal Contingent Negative Variation Drifts During Facial Expression Judgment in Schizophrenia Patients. Front Hum Neurosci, 14, 274. 10.3389/fnhum.2020.00274

Zeidan, J., Fombonne, E., Scorah, J., Ibrahim, A., Durkin, M. S., Saxena, S., Yusuf, A., Shih, A., & Elsabbagh, M. (2022). Global prevalence of autism: A systematic review update. Autism Res, 15(5), 778–790. 10.1002/aur.2696

